# Insights into the bottromycin A_2_ mechanism of action

**DOI:** 10.1101/2025.08.19.671025

**Authors:** Inna A. Volynkina, Aleksandr A. Grachev, Alexei Livenskyi, Daria K. Yagoda, Pavel S. Kasatsky, Olga A. Tolicheva, Ekaterina S. Komarova, Alexey E. Tupikin, Vera A. Alferova, Anastasiia O. Karakchieva, Arina A. Nikandrova, Mikhail V. Biryukov, Yuliya V. Zakalyukina, Lubov V. Dorofeeva, Yuriy A. Ikhalaynen, Igor A. Rodin, Dmitrii A. Lukianov, Marsel R. Kabilov, Alena Paleskava, Andrey L. Konevega, Petr V. Sergiev, Olga A. Dontsova

**Affiliations:** Department of Chemistry, Lomonosov Moscow State University, Moscow, 119234, Russia; Center for Molecular and Cellular Biology, Moscow, 121205, Russia; Molecular and Radiation Biophysics Division, Petersburg Nuclear Physics Institute named by B.P. Konstantinov of National Research Center “Kurchatov Institute”, Gatchina, 188300, Russia; Institute of Gene Biology, Russian Academy of Sciences, Moscow, 119334, Russia; Faculty of Bioengineering and Bioinformatics, Lomonosov Moscow State University, Moscow, 119234, Russia; Belozersky Institute of Physico-Chemical Biology, Lomonosov Moscow State University, Moscow, 119234, Russia; Institute of Chemical Biology and Fundamental Medicine, Siberian Branch of the Russian Academy of Sciences, Novosibirsk, 630090, Russia; Department of Functioning of Living Systems, Shemyakin-Ovchinnikov Institute of Bioorganic Chemistry, Moscow, 117997, Russia; Department of Biology, Lomonosov Moscow State University, Moscow, 119234, Russia; Department of Soil Science, Lomonosov Moscow State University, Moscow, 119234, Russia; All-Russian Collection of Microorganisms (VKM), G.K. Skryabin Institute of Biochemistry and Physiology of Microorganisms, Pushchino Scientific Center for Biological Research, Russian Academy of Sciences, Pushchino, 142290, Russia; Institute of Biomedical Systems and Biotechnologies, St. Petersburg Peter the Great Polytechnic University, Saint Petersburg, 195251, Russia; Centre for Nano, Bio, Info, Cognitive, and Social Sciences and Technologies (NBICS Center), National Research Center “Kurchatov Institute”, Moscow, 123182, Russia; Institute of Functional Genomics, Lomonosov Moscow State University, Moscow, 119234, Russia

**Author notes:** To whom correspondence should be addressed., Correspondence may also be addressed to Andrey L. Konevega., Correspondence may also be addressed to Petr V. Sergiev.

## Abstract

The spread of antimicrobial resistance among pathogenic bacteria poses a threat for modern medicine, highlighting the need for the discovery and development of new potential therapeutic agents. Bottromycin A_2_ (BotA2) represents a promising candidate for future drug development, exhibiting activity against clinically relevant methicillin-resistant *Staphylococcus aureus*, vancomycin-resistant *Enterococcus*, and mycoplasma. However, its exact mechanism of action has not been fully elucidated until now. Here, we demonstrate that BotA2 inhibits bacterial translation showing unique context specificity with regard to the mRNA coding sequence. By using high-throughput toe-printing combined with deep sequencing (toe-seq analysis), we show that BotA2 induces ribosome stalling predominantly when a glycine codon enters the A-site of the ribosome, with stalling efficiency independent of codons located in the P– and E-sites. Our biochemical and biophysical data reveal that BotA2 arrests glycine-containing ternary complexes on the ribosome thereby preventing the full accommodation of incoming Gly-tRNA^Gly^ in the peptidyl transferase center. Altogether, our findings uncover a completely novel, previously undescribed mechanism of translation inhibition based on the context-specific immobilization of ternary complexes on elongating ribosomes.

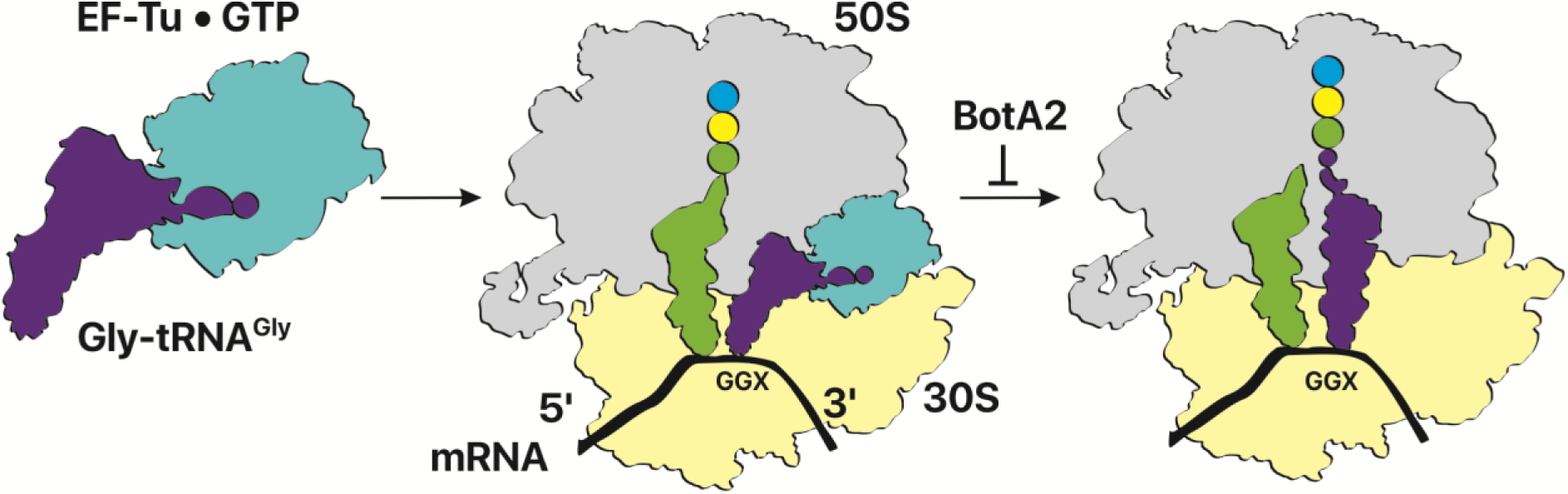
GRAPHICAL ABSTRACT.

## INTRODUCTION

Bottromycin A_2_ (BotA2) is a ribosomally synthesised and post-translationally modified peptide, which was first reported in 1957 [1]. It was purified from the fermentation broth of *Streptomyces bottropensis* and found to be active against Gram-positive bacteria [2] and mycoplasma [3,4]. Later on, other structurally related compounds, named as bottromycin B_2_ [2], C_2_ [5], and D [6] were discovered. Together with BotA2, they constitute a unique class of macrocyclic peptide antibiotics ‒ bottromycins ‒ gaining a special interest nowadays as promising antimicrobial agents [7].

The bottromycins originate from a ribosomally biosynthesized precursor peptide undergoing a series of post-translational modifications catalyzed by enzymes encoded in the bottromycin (*btm*) gene cluster (Figure 1). For a long time, the biosynthesis of bottromycins has been studied step by step [8]. Despite the many published works, we suggest using the *Streptomyces scabies* nomenclature here onwards in order to prevent confusion [9]. A distinctive feature of bottromycins is that their precursor peptide (BtmD) consists of a small core peptide (8 aa residues) bearing a *C*-terminal follower peptide (35-41 aa residues) instead of the canonical *N*-terminal leader sequence [10]. This follower peptide is presumably recognized by biosynthetic enzymes to recruit them to the core peptide, and is eventually removed by proteases upon bottromycin maturation. The proposed biosynthetic pathway starts from the removal of *N*-terminal *N*-formylmethionine (fMet) catalyzed by BtmM aminopeptidase (Figure 1B) [11,12]. Then three radical SAM (*S*-adenosyl-L-methionine) methyltransferases (BtmC, BtmG, BtmK) catalyze *C*-methylation of Pro2, Val4, Val5, and Phe6 residues [13], while two YcaO-domain proteins (BtmE, BtmF) facilitate intramolecular cyclization [14], followed by BtmI-mediated removal of the follower peptide [15]. At the next step, BtmH catalyzes the epimerization of L-Asp7 to D-Asp7 [16], with subsequent oxidative decarboxylation of Cys8 accomplished by the P450 enzyme BtmJ [14,17]. The final step is the *O*-methylation of Asp7, which is catalyzed by the SAM-dependent *O*-methyltransferase BtmB [13,14]. The mature bottromycin is probably exported through the predicted BtmA transporter [11].

**Figure 1.**
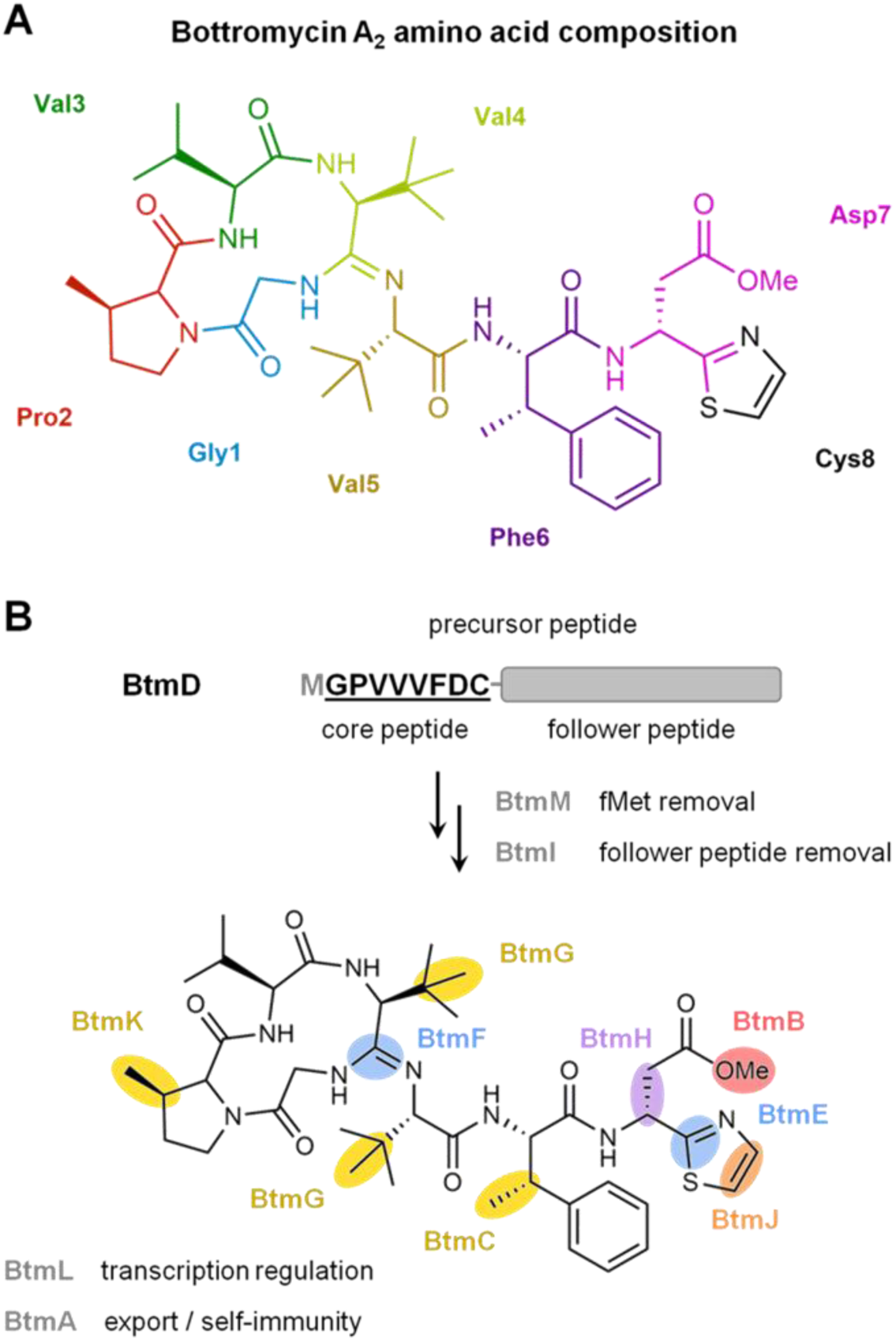
Structure and biosynthesis of bottromycin A_2_. (**A**) Amino acid composition of bottromycin A_2_ (BotA2). Each color highlights individual amino acids of the core peptide, modified during BotA2 maturation. (**B**) Post-translational modifications of the precursor peptide and biosynthetic enzymes responsible for BotA2 maturation. Gene names are given in accordance with the *btm* gene cluster in *Streptomyces scabies* [9].

Despite instability of BotA2 in oral and parenteral administration [4,18], primarily caused by hydrolysis of the methyl ester of Asp7 under physiological conditions (e.g., in blood plasma) [19], BotA2 represents a promising scaffold in drug design, demonstrating activity against a wide range of bacterial strains [5]. Of particular interest is the activity of BotA2 on methicillin-resistant *Staphylococcus aureus* and vancomycin-resistant *Enterococcus* [20,21], as well as its moderate activity against *Mycobacterium* sp. and other *Staphylococcus aureus* clinical isolates resistant to erythromycin, carbomycin, tetracycline, and penicillin [2,5].

Moreover, BotA2 has proven its efficiency at combating mycoplasma infections *in vitro* [3,4,22], being even more potent than classical therapeutic agents (macrolides, tetracyclines) used for the treatment of mycoplasmoses [23]. In addition, BotA2 was also found to be active *in vitro* against the Gram-negative phytopathogenic bacterium *Xanthomonas oryzae* pv. *oryzae*, known to cause bacterial leaf blight of rice [24]. This emphasizes the possibility of BotA2 application in plant protection.

Promising antibacterial activity has prompted researchers to develop chemical derivatives of bottromycin A_2_ in order to obtain more stable compounds [18,19,25–31]. For instance, bottromycin A_2_ hydrazide, reported in 1966, was much more stable in the bloodstream and active *in vivo* against *Mycoplasma gallisepticum* in the infected chickens [4]. However, it demonstrated 8-16 times lower antibacterial activity *in vitro* compared to BotA2 [19]. Over time, the accumulated knowledge allowed characterizing structure-activity relationships of BotA2 and its derivatives, revealing that the ester and thiazole moieties could be altered, while the rest of the molecule is essential for antibacterial properties [30]. To date, only two synthesized BotA2 derivatives ‒ propyl and ethyl ketones replacing the ester group ‒ were shown to have both improved plasma stability and antibacterial activity at least comparable to that of the parental BotA2 [19]. These two compounds, especially the propyl ketone, might be considered as prospective candidates for future drug development.

Although the biosynthesis of bottromycin A_2_ is well studied, its molecular mechanism of action remains obscure. Since 1957, a total of 13 works have been published describing BotA2 mode of action, with the last one being published in 1983 [32]. Overall, these data suggest that BotA2 inhibits translation by targeting the bacterial ribosome [33,34], while the details of its action remain unclear. There have been conflicting reports as to the ability of BotA2 to inhibit the puromycin reaction [35–38], and many proposals that BotA2 exerts its influence either through inhibiting EF-G-dependent translocation [36,39] or through lowering the affinity of aminoacyl-tRNAs (aa-tRNAs) for the A-site [32,38,40].

From all these experimental data, there are several important observations that deserve to be mentioned. In the earliest works, the authors noted that the activity of BotA2 highly depends on the presence of C and G nucleotides in the mRNA template [33,35]. Thus, BotA2 was more efficient at inhibiting *in vitro* translation on poly(C), poly(UC), and poly(UG) mRNAs, compared to poly(A) and poly(U). Moreover, BotA2 was shown not to affect aminoacylation of some tRNAs (tRNA^Leu^, tRNA^Phe^, and tRNA^Pro^) [33]. Non-enzymatic binding of aa-tRNAs (Pro-tRNA^Pro^, Phe-tRNA^Phe^) to ribosomes was also uninfluenced by BotA2 [33,37,41]. Although, BotA2 was able to interfere with the binding of some ternary complexes (aa-tRNA·EF-Tu·GTP) in a concentration-dependent manner, with the inhibition rate being 30-50% [32,40]. Furthermore, BotA2 was demonstrated not to inhibit the puromycin-mediated release of nascent peptides in the *in vivo* system based on *Bacillus megaterium* protoplasts [42]. Likewise, no substantial inhibition of puromycin reaction was observed by using polysomes purified from *Escherichia coli* cells, even when the reactions were supplemented with elongation factor EF-G and GTP [32,43]. Along with this observation, BotA2-induced inhibition of polypeptide synthesis was insensitive to the presence of EF-G·GTP, suggesting that BotA2 is unlikely to directly affect translocation [37,40]. Not surprisingly, BotA2 was shown not to affect EF-G-dependent GTP hydrolysis [37]. Another important finding was that the presence of an excess of 50S over 30S ribosomal subunit decreased the inhibitory effect of BotA2 on polypeptide synthesis, while such an effect was not observed using an excess of 30S over 50S [34]. This indicates that BotA2 presumably interacts with the 50S ribosomal subunit. However, all these preceding studies do not allow us to draw an unambiguous conclusion about the detailed mechanism of action of BotA2.

Herein we present insights into the molecular mechanism of action of BotA2. By using biochemical and biophysical approaches, we uncovered that BotA2 inhibits bacterial translation showing unique context specificity with regard to the mRNA coding sequence. BotA2 was found to induce ribosome stalling exclusively when a Gly codon is in the A-site of the ribosome, with stalling efficiency independent of the codon located in the P-or E-sites. Delving into the details of BotA2 action, we identified that BotA2 does not target Gly-tRNA synthetase and does not provoke mistranslation of Gly codons, rather completely abolishes peptide bond formation with the incoming Gly residue. We were able to show that BotA2 does not interfere with EF-Tu-dependent Gly-tRNA^Gly^ delivery to the ribosome, but rather traps the ternary complex, preventing aa-tRNA from its full accommodation into the ribosomal A-site. Taking into account amino acid specificity of BotA2, we envisage a novel, previously undescribed mechanism of translation inhibition based on specific immobilization of Gly-tRNA^Gly^ in the unaccommodated (likely A/T) state on the ribosome.

## MATERIAL AND METHODS

### Antibiotics

Bottromycin A_2_ (BotA2) and bottromycin A_2_ carboxylic acid (BotCA) standards were kindly provided by Prof. Rolf Müller and Dr. Joy Birkelbach, Helmholtz Institute for Pharmaceutical Research Saarland (HIPS), Helmholtz Centre for Infection Research (HZI), Saarbrücken, Germany. BotA2 and BotCA used in experiments were purified from culture broth of *Streptomyces* sp. VKM Ac-2945^T^ as described below. Mupirocin (Mup) was purchased from pharmacy. Erythromycin (Ery), chloramphenicol (Chl), levofloxacin (Lev), thiostrepton (Ths), borrelidin (Borr), cycloheximide (Chx), kanamycin (Kan), rifampicin (Rif), streptomycin (Str), kirromycin (Kirr) used as reference agents, were purchased from Sigma-Aldrich, Burlington, MA, USA. Tetracenomycin X (TcmX) was purified from culture broth of *Amycolatopsis* sp. A23 as described previously [44]. Microcin C (McC) was purified as decribed in [45].

### Cultivation of the producing strain, fractionation, and isolation of BotA2 and BotCA

The producing strain *Streptomyces* sp. VKM Ac-2945^T^ was provided from the All-Russian Collection of Microorganisms, Pushchino, Russia. It was originally isolated from soil collected in the Belgorod region in 1993. The strain was cultivated in 750 mL Erlenmeyer flasks containing 200-250 mL of soy-glycerol medium [2% soy flour, 1.5% glycerol, 0.2% yeast extract, 0.5% CaCl_2_, 0.1% NaCl, 0.1% K_2_HPO_4_, tap water, pH 7.2] at 28 °C with constant shaking (200 rpm, Innova^®^ 44 Shaker, New Brunswick Scientific, Edison, NJ, USA) until the appearance of pronounced antibacterial activity (∼10-12 days). The activity of fermentation broth and eluted fractions was assessed using the *E. coli lptD^mut^* strain. The culture broth was further separated from the biomass by centrifugation at 20,000 g for 5 min and subjected to solid-phase extraction on LPS-500-H sorbent with 120 μm particle size (“Technosorbent” LLC, Moscow, Russia) using water-acetonitrile mixtures as eluents. Active fractions, containing BotCA (eluted with 30-50% acetonitrile) and BotA2 (eluted with 75-100% acetonitrile) were collected and subjected to preparative RP-HPLC.

Preparative RP-HPLC was performed on a puriFlash^®^ PF-4250 preparative chromatograph (Interchim, Montluçon, France) equipped with a VDSpher 100 C18-E (10 µm, 20×250 mm) column (VDS optilab, Berlin, Germany). UV-Vis detection was carried out at 205 and 254 nm. Eluent A consisted of 0.1% trifluoroacetic acid (TFA) (HPLC grade, Fisher Scientific, Loughborough, UK) in ultrapure water; eluent B was acetonitrile (HPLC gradient grade, Fisher Scientific, Loughborough, UK) containing 0.1% TFA. Ultrapure water was obtained using the HyperPureX purification system (Hyperpurex Instrument Technology, Shanghai, China). The flow rate was maintained at 20 mL/min. For BotCA, the gradient program was 0–25–28–33 min, 29–29–95–95% B, with a retention time of 24.5 min. For BotA2, the program was 0–15– 25–30 min, 30–40–95–95% B, with a retention time of 19.3 min (Supplementary Figure S1).

The identity of the isolated compounds was first confirmed by analytical RP-HPLC using BotA2 and BotCA standards, and then by high-resolution mass spectroscopy (HRMS) analysis. Analytical RP-HPLC was carried out on an Agilent 1100 system (Agilent Technologies, Santa Clara, CA, USA) with a quaternary pump, equipped with an Ultisil XB-C18 (5 µm, 4.6×250 mm) column (Welch Materials, Shanghai, China) and a corresponding Ultisil XB-C18 (5 µm, 4.6×20 mm) guard column (Welch Materials, Shanghai, China). Eluent A was ultrapure water, eluent B was acetonitrile, and eluent C was 2% aqueous TFA. For BotA2, the analytical method applied was 0–10–12–14 min, 40–70–95–95% B, with a constant addition of 5% C, yielding a retention time of 6.9 min. For BotCA, the method was 0–30–31–34 min, 29–29–95–95% B, with a constant addition of 5% C, yielding a retention time of 22.6 min. The identity of the samples was further confirmed by LC-HRMS/MS analysis using the Orbitrap Exploris 120 mass spectrometer combined with the Vanquish UHPLC system (Thermo Fisher Scientific, Waltham, MA, USA), equipped with a reversed-phase Acclaim™ 120 C18 (2.2 µm, 2.1×150 mm) column (Dionex, Sunnyvale, CA, USA). Eluent A was 0.1% formic acid in ultrapure water, eluent B was 0.1% formic acid in acetonitrile. Chromatographic separation was carried out in gradient elution mode at a flow rate of 0.4 mL/min. Gradient parameters were as follows: 0–5 min, 5% B; 5–15 min, 5–95% B; 15–18 min, 95% B. The injection volume was 20 µL, the column oven temperature was 40°C, and the autosampler temperature was 4°C. Heated electrospray ionization (ESI) source parameters were as follows: positive (or negative) ionization; ion transfer voltage—3,500 (2,500) V; sheath/auxiliary/sweep gas— 50/10/1 arb; ion transfer temperature—325 °C; vaporizer temperature—350 °C. Mass spectra were recorded with a full scan from 50 to 1500 m/z. The resolution was 30,000 (for both MS^1^ and MS^2^ spectra), the RF lens—70%, the number of microscans—3, and the data type— centroid. Higher-energy collisional dissociation (HCD) fragmentation parameters were as follows: isolation window—1 m/z with isolation offset turned off; collision energy—15, 30, 45, 60 V. For BotCA, the observed ions were [M−H]^−^ at m/z 807.4224 and [M+H]^+^ at m/z 809.4366 (calculated for C_41_H_60_N_8_O_7_S: [M−H]^−^ 807.4227, Δ 0.4 ppm; [M+H]^+^ 809.4384, Δ 2 ppm). For BotA2, the observed ions were [M−H]^−^ at m/z 821.4382 and [M+H]^+^ at m/z 823.4528 (calculated for C_42_H_62_N_8_O_7_S: [M−H]^−^ 821.4384, Δ 0.2 ppm; [M+H]^+^ 823.4540, Δ 1.5 ppm). The MS^2^ fragmentation patterns of both compounds are provided in Supplementary Figure S2 and display all the characteristic fragment ions.

### Bacterial strains and cultivation conditions

The *Escherichia coli* BW25113 strain with a partial deletion of the *lptD* gene, codons 330 to 352 (referred to here as *E. coli lptD^mut^*) was kindly provided by Dr. Alexander S. Mankin, University of Illinois, Chicago, IL, USA [46]. The *E. coli* JW5503-KanS strain carrying a deletion of the *tolC* gene (referred to here as *E. coli ΔtolC*) was obtained as described in [47] from the *E. coli* JW5503 strain by removing the kanamycin resistance cassette. *E. coli* JW5503 and *E. coli* BW25113 were kindly provided by Prof. Hironori Niki, National Institute of Genetics, Mishima, Shizuoka, Japan [48]. *E. coli ΔtolC* cells were transformed with pJC27 plasmids, encoding either a native or mutant *lacZ* gene [49]. The following mutations at the catalytic residue of β –galactosidase were used: Glu537Asp (GAA→GAC), Glu537Gly (GAA→GGA/GGG), corresponding to the plasmids pJC27-GAC, pJC27-GGA, and pJC27-GGG.

The *E. coli* SQ110 strain with the deletion of six rRNA operons (except for *rrnE*) and the *tolC* gene (referred to here as SQ110 ΔtolC) was kindly provided by Dr. Alexander S. Mankin [46]. Mutants resistant to Chl, Ery, or both of them were selected using the SQ110 ΔtolC strain, according to the procedure reported previously [44]. The *E. coli* SQ171 strain with the deletion of all seven rRNA operons and the *tolC* gene (referred to here as SQ171 ΔtolC), and transformed with the pCSacB (Kan^R^) plasmid, which contains the wild-type *rrnB* operon, was kindly provided by Dr. Alexander S. Mankin [50,51]. The pCSacB (Kan^R^) plasmid was further replaced with the pLK35 (Amp^R^) or pAM552 (Amp^R^) plasmid encoding either the native (WT) or mutated *rrnB* operon, according to the procedure described previously [44].

*Bacillus subtilis* 168 was kindly provided by Vera A. Alferova, Shemyakin-Ovchinnikov Institute of Bioorganic Chemistry, Moscow, Russia. *B. subtilis* 168 pHT01-cat and *B. subtilis* 168 pHT01-cfr were obtained as described previously [52].

Unless otherwise stated, all aforementioned bacterial strains were grown at 37 °C in Miller’s Lysogeny Broth (LB) medium supplemented with antibiotics, if required. *E. coli lptD^mut^* and *E. coli ΔtolC* transformed with the pJC27 plasmids were cultivated in the presence of 10 μg/mL chloramphenicol. SQ110 ΔtolC strains were cultivated in the presence of 50 μg/mL kanamycin. SQ171 ΔtolC pLK35 and SQ171 ΔtolC pAM552 strains were cultivated in the presence of 100 μg/mL ampicillin. *B. subtilis* 168 pHT01-cat was cultivated in the presence of 5 μg/mL chloramphenicol. *B. subtilis* 168 pHT01-cfr was cultivated in the presence of 3 μg/mL chloramphenicol. Other bacterial strains were cultivated without antibiotics.

### Agar diffusion assay on a panel of resistant mutants

Overnight cultures of SQ110 ΔtolC and SQ171 ΔtolC cells grown in the presence of relevant antibiotics were plated on 1.5% LB-agar solid medium supplied either with 50 μg/mL kanamycin or with 100 μg/mL ampicillin, respectively, and left to dry for a while until the next step. The following antibiotics were applied on the surface of agar plates coated with SQ110 ΔtolC or SQ171 ΔtolC cells: erythromycin (Ery, 5 mg/mL), levofloxacin (Lev, 25 μg/mL), tetracenomycin X (TcmX, 5 mg/mL), chloramphenicol (Chl, 2 mg/mL). In the center of each agar plate, a well was made and filled with 50 μL of 1 mM BotA2 dissolved in 30% acetonitrile aqueous solution. The plates were incubated overnight at 37 °C and scanned with the ChemiDoc™ Imaging System (Bio-Rad Laboratories, Hercules, CA, USA) using the “Cy5-blot” channel (emission filter 695 ± 50 nm). Images were analysed and visualized using the Image Lab™ software (version 6.0.1, Bio-Rad Laboratories, Hercules, CA, USA).

### Determination of minimum inhibitory concentration (MIC)

MIC values were determined using the liquid broth microdilution assay in 96-well sterile plates and a total volume of 100 μL per well, as described in [53]. Antibiotics were dissolved in 50% dimethyl sulfoxide (DMSO) aqueous solution to obtain 20 mM stock solutions. Then they were diluted 1:50 with 100 μL of fresh LB medium. Subsequently, 2-fold serial dilutions were performed and bacterial cells in the log growth phase diluted 1:100 with LB medium were added to each well. The plates were incubated overnight (16-20 h) at 37 °C with constant shaking at 200 rpm (Shaker-Incubator ES-20/80, BioSan, Riga, Latvia). Cell growth was assessed by scanning the OD_600_ with the VICTOR X5 Multilabel Plate Reader (PerkinElmer, Waltham, MA, USA). Minimum inhibitory concentration was defined as the lowest concentration of an examined compound at which the growth of the bacterial strain was completely inhibited. For each MIC measurement, at least two biological replicates were performed.

### *In vitro* translation assays

*In vitro translation in bacterial cell-free systems.* Firefly luciferase (Fluc) mRNA was prepared using the MEGAscript™ T7 Transcription Kit (Thermo Fisher Scientific, Waltham, MA, USA). Translation reactions (3 μL total volume) were carried out with the PURExpress^®^ In vitro Protein Synthesis Kit (New England BioLabs, Ipswich, MA, USA) according to the manufacturer’s protocol. Alternatively, the reactions (5 μL total volume) were carried out with *E. coli* S30 Extract System for Linear Templates (Promega, Madison, WI, USA) according to the manufacturer’s instructions. Each reaction was supplied with 2 U of RiboLock RNase Inhibitor (Thermo Fisher Scientific, Waltham, MA, USA), 0.17 mM of D-luciferin sodium salt (Sigma-Aldrich, Burlington, MA, USA), 60 ng of Fluc mRNA, and a chemical compound or nuclease-free water instead. Before the addition of mRNA, reaction tubes were pre-incubated at room temperature (RT) for 5 min and then placed back on ice. After the addition of mRNA, reaction mixtures were transferred into the 384-well white microplate. Chemiluminescence was continuously measured using the VICTOR X5 Multilabel Plate Reader (PerkinElmer, Waltham, MA, USA) at 37 °C for 1 h. The maximum Fluc accumulation rates were calculated (as an increment of light intensity units per second) and defined as the efficiency of translation reactions. The values were normalized to a positive control (nuclease-free water, assigned a value of 100%). The IC_50_ values of translation inhibition were calculated and visualized using the GraphPad Prism software (version 10.1.2, Dotmatics, Boston, MA, USA).

*In vitro translation in a mammalian cell-free system.* The whole cell extract was prepared from the HEK293T cell line as described previously [54] with minor modifications: harvested cells were not treated with lysolecithin buffer. Instead, they were immediately suspended in an equal volume of ice-cold hypotonic extraction buffer [20 mM HEPES-KOH (pH 7.5), 10 mM KOAc, 1 mM Mg(OAc)_2_, 4 mM DTT and cOmplete™ EDTA-free Protease Inhibitor Cocktail (Roche, Basel, Switzerland)], incubated for 5 min on ice and disrupted in a tiny Dounce homogenizer by 20-25 strokes. The lysate was clarified by centrifugation for 10 min at 10,000 g at 4 °C. Aliquots were frozen in liquid nitrogen and stored at –80 °C.

Translation reactions were performed as described previously [55,56]. Briefly, each reaction was carried out in a total volume of 10 μL and contained 5 μL of the HEK293T whole cell extract, translation buffer [20 mM HEPES-KOH (pH 7.5), 8 mM creatine phosphate, 1 mM DTT, 0.5 mM spermidine-HCl, 1 mM ATP, 0.2 mM GTP, 100 mM KOAc, 1 mM Mg(OAc)_2_ and 25 μM of each amino acid], 2 U of Creatine Phosphokinase from rabbit muscle (Sigma-Aldrich, Burlington, MA, USA), 2 U of RiboLock RNase Inhibitor (Thermo Fisher Scientific, Waltham, MA, USA), 0.5 mM of D-luciferin sodium salt (Sigma-Aldrich, Burlington, MA, USA), 1 μL of either an antibiotic solution or nuclease-free water as a positive control, and 100 ng of m^7^G-capped and polyadenylated Fluc mRNA with the β-actin 5’ UTR. The latter one was prepared using the mMESSAGE mMACHINE™ T7 Transcription Kit (Thermo Fisher Scientific, Waltham, MA, USA) and added as 1 μL water solution to the translation mixtures after they were pre-incubated for 5 min at 30 °C. Then, the mixtures were transferred into the pre-warmed 384-well white microplate, covered with a PCR plate seal and incubated in the CLARIOstar^®^ Plus Microplate Reader (BMG Labtech, Ortenberg, Germany) at 30 °C with continuous measurement of the luciferase activity. The maximum Fluc accumulation rates were calculated and defined as the efficiency of translation reactions. The values were normalized to a positive control (assigned a value of 100%) and visualized using the GraphPad Prism software (version 10.1.2, Dotmatics, Boston, MA, USA).

### Toe-printing assay

The toe-printing analysis of drug-dependent ribosome stalling was performed essentially as previously described [57] with some minor modifications indicated below. Linear DNA templates RST1 and RST3 (Supplementary Table S2) were generated by PCR using the following pairs of partially complementary primers – RST1-fwd + RST1-rev, and RST3-fwd + RST3-rev, respectively (Supplementary Table S1). DNA templates containing short ORFs (Supplementary Table S2) were generated by PCR using the pRFPCER plasmid [58] as a template, the CER-FAM-R reverse primer (Supplementary Table S1), and a set of different forward primers, listed in the Supplementary Table S1. MG template was generated using the M-GGC forward primer and NEW_CER_FAM reverse primer instead of CER-FAM-R. The resulting templates (0.3 pmol) were expressed in a cell-free bacterial transcription-translation coupled system using the PURExpress^®^ In Vitro Protein Synthesis Kit (New England BioLabs, Ipswich, MA, USA) in a total volume of 5 μL, following the manufacturer’s protocol. All antibiotics were added to the final concentration of 50 μM. Negative control samples (indicated as “−”) were supplemented with nuclease-free water instead of antibiotics. Before the addition of a DNA template, reaction mixtures were pre-incubated at RT for 5 min.

Following the addition of a DNA template, the reactions were incubated at 37 °C for 20 min, then 1 pmol of the [^32^P]-labeled NV1 or FAMnoFAM primer (Supplementary Table S1) and 2 U of the AMV Reverse Transcriptase (Roche, Basel, Switzerland) were added, and reaction tubes were additionally incubated at 37 °C for 15 min. Reactions were terminated by the addition of 1 μL of 10 M NaOH followed by incubation at 37 °C for 15 min. The pH was further neutralized and stabilized by the addition of an equivalent amount of 10 N HCl, and then 200 μL of Resuspension Buffer [0.3 M NaOAc (pH 5.5), 5 mM EDTA, 0.5% SDS].

Thereafter, the samples were subjected to cDNA fragments purification using the QIAquick^®^ PCR Purification Kit (Qiagen, Hilden, Germany) according to the manufacturer’s protocol, instead of phenol-chloroform extraction. Resulting cDNA solutions were dried at 95 °C and dissolved in 10 μL of Formamide Loading Buffer [98% formamide, 10 mM EDTA (pH 8.0), 0.1% bromophenol blue, 0.1% xylene cyanol]. The sequencing reactions were performed using the USB^®^ Thermo Sequenase Cycle Sequencing Kit (Affymetrix, Santa Clara, CA, USA) according to the manufacturer’s protocol. The cDNA fragments, along with sequencing reactions, were resolved on a 6% polyacrylamide sequencing gel containing 7 M urea in TBE buffer [90 mM Tris base, 90 mM boric acid, 2 mM EDTA, pH 8.3]. After electrophoresis, the gel was dried using the Savant™ Universal Vacuum System Plus with VaporNet^®^ (UVS400A, Thermo Fisher Scientific, Waltham, MA, USA) at 80 °C for 1 h, and exposed to the phosphorimager screen overnight. The screen was scanned using the Typhoon™ FLA 9500 Biomolecular Imager (GE Healthcare, Chicago, IL, USA) or the FujiFilm Fluorescent Image Analyzer FLA-3000 (Fujifilm, Tokyo, Japan), and the images were processed using the Image Lab™ software (version 6.0.1, Bio-Rad Laboratories, Hercules, CA, USA).

### RelE-printing assay

The RelE-printing analysis was performed in a similar way as described above for the toe-printing, but with some modifications indicated below. The RelE protein (0.8 mg/mL stock solution), purified as described in [59], was kindly provided by Dr. Dmitry E. Andreev, Belozersky Institute of Physico-Chemical Biology, Lomonosov Moscow State University, Moscow, Russia. RelE was diluted 1:10 in Pure System Buffer [9 mM Mg(OAc)_2_, 5 mM KH_2_PO_4_, 95 mM potassium glutamate, 5 mM NH_4_Cl, 0.5 mM CaCl_2_, 1 mM spermidine, 8 mM 1,4-diaminobutane, 1 mM DTT, pH 7.3] prior the experiment. For control reactions, nuclease-free water was diluted 1:10 in Pure System Buffer. Following the incubation of translation reactions at 37 °C for 15 min, 0.5 μL of the RelE solution or the control solution was added to each sample and the incubation was continued at 37 °C for 15 min. Next, 1 pmol of the [^32^P]-labeled FAMnoFAM primer (Supplementary Table S1) and 2 U of the AMV Reverse Transcriptase (Roche, Basel, Switzerland) were added, and reaction tubes were additionally incubated at 37 °C for 15 min. The subsequent experimental procedure was exactly the same as described above for the toe-printing assay.

### Toe-seq assay

*Preparation of samples for toe-seq.* The toe-seq analysis of BotA2 context specificity was performed according to the procedure reported in [60]. In brief, a library of linear DNA templates was generated by PCR in the preceding work. One template consisted of a T7 promoter, a 5’ UTR (one of three types), an ORF containing a start codon, 30 randomized nucleotides, and 7 constant codons, followed by a 3’ UTR (one of two types) and a barcode region. Next generation sequencing (NGS) of the library resulted in a dictionary representing each barcode matched up with its ORF sequence and both UTR sequences. The library of DNA templates was expressed in a bacterial transcription-translation coupled system using the PURExpress^®^ In Vitro Protein Synthesis Kit (New England BioLabs, Ipswich, MA, USA) in the presence of 50 μM BotA2. Reactions supplied with nuclease-free water instead of BotA2 served as untreated controls. All reactions were prepared in two biological replicates. AMV Reverse Transcriptase (Roche, Basel, Switzerland) was further applied to synthesize cDNA fragments, which were further purified and elongated at their 3’ end by 5 U of Terminal Transferase (New England BioLabs, Ipswich, MA, USA) supplied with 4 μM dATP. Poly(A)-tailed cDNA was then converted into dsDNA by Klenow Fragment (3’→5’ exo-) (New England BioLabs, Ipswich, MA, USA) according to the manufacturer’s protocol. After that, dsDNA fragments were additionally amplified with high-fidelity PCR. Purified PCR products were subjected to library preparation using the MGIEasy Universal DNA Library Prep Set (MGI Tech, Shenzhen, China) and sequenced on the MGIseq-2000 (MGI Tech, Shenzhen, China) at the Genomics Core Facility (ICBFM SB RAS, Novosibirsk, Russia).

*Computational processing of toe-seq data.* Raw FastQ sequencing data were processed according to the procedure reported in [60]. All the computations described below were performed using in-house Python scripts and the previously obtained dictionary. Parameters of the resulting toe-seq datasets are given in the Supplementary Table S3. For each mRNA, read counts per nucleotide position were calculated as the number of reads ending at each position. Reads terminated at 5’ UTR were summed and designated as FL values (the quantity of full-length cDNA fragments). Thereafter read counts per position were normalized to the total number of mapped reads and then multiplied by 10^6^, resulting in the so-called counts per million (CPM). For each sense codon, normalized read counts associated with positions +16, +17, and +18 relative to the first nucleotide of the codon were summed up and divided by the total number of CPM of a given mRNA (*CPM_mRNA_*). The resulting values reflect ribosome density distribution along the coding sequence and correspond to codons located in the P-site of the ribosome. In addition, all CPM values associated with positions located between the start codon and the start codon +15 nucleotides were summed with the previously calculated FL values and divided by the corresponding *CPM_mRNA_*. Next, we normalized the data of antibiotic-treated samples to that of control samples (nuclease-free water) by subtracting the corresponding ribosome densities. This step additionally allows us to exclude non-specific reverse transcriptase products from further consideration. The resulting positive values represent the distribution of ribosome stalling probability (*StopProbability*) along the ORF sequence, whilst the negative values should be interpreted as truly “impossible” sites for antibiotic-specific ribosome stalling, not merely “zero probability”. The serial number of the codon associated with the maximum probability of ribosome stalling (*MaxStopProbability*) was defined as the stop position (*StopPosition*) at a given mRNA. In the case of mRNAs containing a premature stop codon, only appropriate *StopProbability* values were considered to determine *MaxStopProbability* and *StopPosition*. All *StopProbability* values that fall outside the real range of ORF were excluded from the analysis in advance. Since the accuracy of the *MaxStopProbability* score depends on the mRNA coverage (designated here as *CPM_mRNA_*), we introduced a *Likelihood* metric that takes into account the reliability of the calculated *MaxStopProbability* over the entire dataset. This metric was defined as follows:

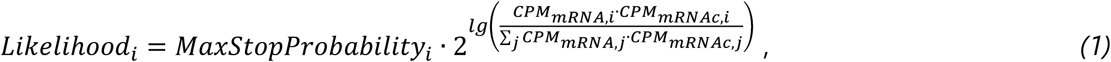

where *CPM_mRNA,i_* and *CPM_mRNAc,i_* stand for the total number of CPM attributed to a given mRNA (with the barcode =) and obtained from antibiotic-treated and control samples, respectively. It is worth noting that we use *Likelihood* as a relative rather than an absolute metric. Likewise, we introduced another parameter named *cpmRNAratio*, which represents the similarity in mRNA coverage between two samples and is calculated as follows:

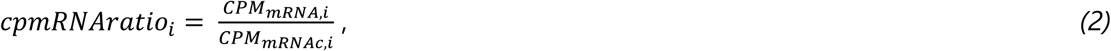

The following criteria were further applied to extract the most relevant ribosome stalling sites from a dataset: (i) *MaxStopProbability* score should be higher than 0, (ii) *cpmRNAratio* should be within the interquartile range (IQR), (iii) mRNAs with *CPM_ORF_*, *CPM_mRNAc_*, and *Likelihood* values below the corresponding 25^th^ percentile should be filtered out. Here, *CPM_ORF_* is the number of CPM in mRNA coding region. The resulting datasets were used for further analysis.

*Analysis of amino acid enrichment at the sites of BotA2-induced ribosome stalling.* Both filtered datasets derived from BotA2-treated samples were used to extract ribosome stalling sites defined by the calculated *StopPositions*. The sequence of each site included amino acid residues corresponding to the codons located in the E-, P-, and A-sites of arrested ribosomes. Sites associated with ribosome stalling at the start and stop codons, as well as at the second codon and codons 11-17, were excluded from the analysis to consider only variable region of mRNA. A total of 12,245 sites were applied as foreground sequences to the pLogo tool [61]. To generate background sequences, we performed the same calculations with respect to control datasets. However, we omitted the step of ribosome density subtraction and used a limited set of filtration criteria (only points (i) and (iii)). Apparent ribosome stalling sites for control samples were extracted in exactly the same way as for BotA2-treated samples and applied as background sequences to the pLogo tool.

*Analysis of codon enrichment at the sites of BotA2-induced ribosome stalling.* Each filtered dataset derived from BotA2-treated samples was used to calculate relative occurrence of individual codons at E-, P-, and A-site positions among the identified stalling sites. Only sites associated with *StopPositions* 3-10 were included in the analysis, as they correspond to the variable region of mRNA. Likewise, relative occurrence of individual codons was calculated for apparent ribosome stalling sites from control datasets. The resulting values for control datasets were further subtracted from the corresponding values for the BotA2-treated samples to account for possible codon bias. Therefore, the resulting negative values conform to underrepresented codons, while positive values indicate overrepresented ones.

### Metabolic labeling assay

Overnight culture of *E. coli lptD^mut^* was diluted to an OD_600_ of 0.1 in MOPS minimal medium containing 0.4% glycerol without antibiotics and grown at 37 °C for 1 h. Then, 50 μL of [^32^P]-orthophosphoric acid (1 mCi/mL, 8500 Ci/mmole) was added to 1 mL of *E. coli* culture to achieve a final radioactivity of 50 μCi/mL, followed by incubation at 37°C for 50 min.

Bottromicyn A_2_, microcin C and mupirocin were added to *E. coli* cells and incubation continued for 30 min at 37 °C. Final concentrations of antibiotics were 20 μM for bottromycin A_2_, 20 μM for microcin C, and 60 μM for mupirocin, which exceed the corresponding MICs.

Metabolically radiolabeled nucleotides were extracted with formic acid and subjected to thin-layer chromatography as described in [62], for subsequent detection by autoradiography.

### Global analysis of tRNA aminoacylation level (GATRAL)

Overnight culture of *E. coli lptD^mut^* was diluted 1:50 in fresh LB medium without antibiotics and grown at 37 °C until an OD_600_ of 0.4 was reached. Bottromycin A_2_, microcin C, or no additive (control) was added to *E. coli* cells, followed by an additional incubation at 37 °C for 30 min. Final concentrations of antibiotics were 20 μM for bottromycin A_2_ and 20 μM for microcin C, which exceed the corresponding MICs. Then, tRNA fractions were isolated from 5 mL of culture using the Total RNA and Small RNA Isolation Kit (Biolabmix, Novosibirsk, Russia), according to the manufacturer’s protocol. Aminoacyl-tRNAs were acetylated with acetic anhydride, hydrolyzed via RNase digestion, and analyzed by LC-QTOF-MS, according to the published procedure [63].

LC-QTOF-MS analysis was performed using the Acquity UPLC System (Waters, Milford, MA, USA) coupled to a quadrupole time-of-flight (QTOF) mass spectrometer maXis Impact II (Bruker Daltonics, Bremen, Germany), equipped with an electrospray ionization (ESI) source. The sample (10 μL) was completely dissolved in 20 μL of deionized water. Then, 10 μL solutions were injected into the LC-QTOF-MS system. Chromatographic separation was carried out at 40 °C using a binary gradient at a flow rate of 0.2 mL/min on an Acquity UPLC BEH C18 (1.7 μm, 2.1×100 mm) column (Waters, Milford, MA, USA) coupled with Acquity UPLC BEH C18 VanGuard (1.7 μm 2.1×5 mm) pre-column (Waters, Milford, MA, USA). The mobile phase consisted of 5 mM (NH_4_)OAc (pH 5.2) in water (solvent A), and 5 mM (NH_4_)OAc (pH 5.2) in 90% acetonitrile (MeCN) (solvent B). Stepwise linear gradient: 0-3 min with 2% MeCN, 3-28 min from 2% to 40% MeCN, 28-30 min from 40% to 80% MeCN, 30-35 min with 80% MeCN, 35-37 min from 80% to 2% MeCN, 37-40 min with 2% MeCN. The mass spectrometer was operated in positive ionization mode. Total ion chromatograms (TICs) were acquired using the following settings: nebulizer pressure of 1.8 bar, drying gas flow rate of 5 L/min, and drying gas temperature of 180 °C. Mass spectrometer was calibrated daily using the external calibration standard ES Tuning Mix (Agilent Technologies, Santa Clara, CA, USA). Mass spectra were analyzed using the Compass DataAnalysis software (version 4.3, Bruker Daltonics, Bremen, Germany).

### Agar diffusion assay using misreading error reporter systems

*E. coli ΔtolC* cells transformed with the pJC27 reporter systems were used as described previously [64]. In brief, the overnight culture of *E. coli ΔtolC* pJC27 was diluted 1:8 in 4 mL of warm (50 °C) 0.6% LB-agar supplemented with 10 μg/mL chloramphenicol. After a brief mixing, the cell suspension was poured on top of a 1.5% LB-agar plate (9 cm Petri dish) supplemented with 10 μg/mL chloramphenicol, and 80 μg/ml X-Gal (5-bromo-4-chloro-3-indolyl β-D-galactopyranoside). After the soft agar had solidified, antibiotics were applied on the surface of agar plates containing a reporter strain. The plates were incubated overnight at 37 °C and photographed on camera. The following stocks of antibiotics used were: kanamycin (Kan, 5 mg/mL), rifampicin (Rif, 10 mg/mL), streptomycin (Str, 5 mg/mL), bottromycin A_2_ carboxylic acid (BotCA, 20 mM), bottromycin A_2_ (BotA2, 20 mM), and bottromycin A_2_ (indicated as (●), 500 μM).

### Sample preparation for self-assembled *in vitro* translation

*In vitro* translation system on the basis of individually purified components was assembled in the buffer TAKM_7_ [50 mM Tris-HCl (pH 7.5), 70 mM NH_4_Cl, 30 mM KCl, and 7 mM MgCl_2_] essentially as described in [65] with some modifications indicated below. MG (AUG-GGC), MF (AUG-UUU), and MV (AUG-GUU) mRNAs were obtained by T7 transcription followed by purification on a HiTrap^®^ Q HP anion exchange column (Cytiva, Marlborough, MA, USA) [66]. DNA templates for *in vitro* transcription were amplified by PCR using the following pairs of primers ‒ T7-fwd-1 + FAMnoFAM (for MG), and T7-fwd-2 + LP-rev (for MF and MV). DNA sequences of primers and templates are provided in Supplementary Tables S1 and S2, respectively.

Individual tRNA^Gly^ was prepared as described in [67] with some modifications. Briefly, total *E. coli* tRNA was aminoacylated with 0.5 mM glycine at 37 °C for 30 min using 5% S100 extract. Aminoacyl-tRNA was supplemented with two-fold excess of EF-Tu·GTP to form a ternary complex (aa-tRNA·EF-Tu·GTP) at 37 °C for 5 min. Ternary complex was purified on 1 mL column Protino^®^ Ni-IDA (Macherey-Nagel, Düren, Germany) according to the manufacturer’s guidance. The eluted ternary complex was dissociated by addition of potassium acetate (pH 5.0) to a final concentration of 0.2 M, and aminoacyl-tRNA was phenol-extracted and precipitated with ethanol. The tRNA pellet was dissolved in ddH_2_O and stored at –80 °C. Fluorescently labeled tRNA^Gly^(Prf16/17/20) was prepared as described previously [68].

For a typical reaction of initiation complex formation, 1 µM 70S ribosomes were incubated with 5 µM mRNA, 2 µM fMet-tRNA^fMet^ or 1.5 µM BODIPY-Met-tRNA^fMet^,1.5 µM of IF1, IF2 and IF3 in the TAKM_7_ buffer supplemented with 1 mM GTP and 2 mM DTT at 37 °C for 1 h.

Ternary complexes of aa-tRNA·EF-Tu·GTP were prepared by pre-incubation of EF-Tu with 1 mM GTP, 3 mM phosphoenolpyruvate, 2 mM DTT,1% pyruvate kinase in the TAKM_7_ buffer at 37 °C for 15 min, followed by addition of aa-tRNA and incubation for 5 min. fMet-tRNA^fMet^, BODIPY-Met-tRNA^fMet^, Phe-tRNA^Phe^ and Val-tRNA^Val^ were purified by RP-HPLC and stored at –80 °C. Gly-tRNA^Gly^(Prf16/17/20) was prepared immediately before ternary complex formation. For this, 10 µM tRNA^Gly^(Prf16/17/20) was incubated with 0.2 mM Gly, 3 mM ATP, 2 mM DTT, 0.75 µM Gly-tRNA synthetase in the TAKM_7_ buffer at 37 °C for 40 min. Where indicated, ternary complexes were additionally incubated with bottromycin A_2_ or kirromycin at 37 °C for 5 min prior to experiment. Final concentrations of antibiotics after mixing initiation complex with ternary complex were 100 μM and 150 μM for bottromycin A_2_ and kirromycin, respectively.

### *In vitro* synthesis of fluorescently labeled peptides

*In vitro* synthesis of short fluorescently labeled peptides was performed essentially as described in [66,69] with some modifications indicated below. Linear DNA templates (0.3 pmol) were expressed in a cell-free bacterial transcription-translation coupled system using the PURExpress^®^ Δ(aa, tRNA) Kit (New England BioLabs, Ipswich, MA, USA) in a total volume of 5 μL, according to the manufacturer’s guidance. Final concentrations of reagents in translation mixtures were corrected for the sake of better representation as indicated below. Each reaction consisted of 0.8 μL of Solution A (minus aa, tRNA, from the kit), 1.2 μL of Solution B (from the kit), 0.53 μM *E. coli* Ribosomes (New England BioLabs, Ipswich, MA, USA), 0.16 μM BODIPY-Met-tRNA^fMet^, 0.2 μL of uncharged tRNA (7 mg/mL), 0.3 mM L-serine (Sigma-Aldrich, Burlington, MA, USA), 0.3 mM L-glycine (Sigma-Aldrich, Burlington, MA, USA), 2 U of RiboLock RNase Inhibitor (Thermo Fisher Scientific, Waltham, MA, USA), 15 ng of a DNA template, and 50 μM bottromycin A_2_. Negative control samples (indicated as “−”) were supplemented with nuclease-free water instead of bottromycin A_2_. In parallel, control reactions were assembled to synthesize the following fluorescently labeled peptides *in vitro* – BODIPY-Met, BODIPY-Met-Ser, BODIPY-Met-Ser-Gly, and BODIPY-Met-Ser-Gly-Phe. Control reactions consisted of 1 μL of Solution A (minus aa, tRNA, from the kit), 1.3 μL of Solution B (from the kit), 0.08 μM BODIPY-Met-tRNA^fMet^, 0.35 μL of uncharged tRNA (7 mg/mL), 0.3 mM L-serine (Sigma-Aldrich, Burlington, MA, USA), 0.3 mM L-glycine (Sigma-Aldrich, Burlington, MA, USA), 0.3 mM L-phenylalanine (Sigma-Aldrich, Burlington, MA, USA), 2 U of RiboLock RNase Inhibitor (Thermo Fisher Scientific, Waltham, MA, USA), and 10 ng of a DNA template (M-AGT, M-AGT-GGC, or M-AGT-GGC-F, Supplementary Table S2). The “BODIPY-Met” sample was supplemented with nuclease-free water instead of a DNA template. Before the addition of templates, all reaction tubes were pre-incubated at RT for 5 min and then placed back on ice. Following the addition of templates, reaction mixtures were incubated at 37 °C for 30 min. For synthesis of short fluorescently labeled peptides using the self-assembled *in vitro* translation system, initiation complexes were programmed with either of the three mRNAs ‒ MG (AUG-GGC), MF (AUG-UUU), and MV (AUG-GUU) ‒ and assembled with BODIPY-Met-tRNA^fMet^ in the P-site. Ternary complexes were assembled with the following tRNAs ‒ Gly-tRNA^Gly^, Phe-tRNA^Phe^, and Val-tRNA^Val^. Translation was initiated by adding 0.2 µM ternary complex to 0.1 µM initiation complex followed by incubation at 37 °C for 2 min. Translation reactions were terminated by the addition of 0.29 M NaHCO_3_ followed by incubation at 37 °C for 20 min. Then, the samples were mixed with an equal volume of Formamide Loading Dye [98% formamide, 10 mM EDTA (pH 8.0), 0.1% bromophenol blue], incubated for 3 min at 70 °С, and resolved in a 20×20 cm 12% PAAG containing 7 M urea in TBE buffer [90 mM Tris base, 90 mM boric acid, 2 mM EDTA, pH 8.3]. The gel was scanned using the Typhoon™ FLA 9500 Biomolecular Imager (GE Healthcare, Chicago, IL, USA). Laser of 473 nm (blue LD laser) was used for excitation and DBR1 filter of 530 ± 20 nm was used for emission. Images were processed and visualized using the Image Lab™ software (version 6.0.1, Bio-Rad Laboratories, Hercules, CA, USA).

### Preparation of uncharged tRNA

Total tRNA was isolated from *E. coli* BW25113 culture using the Total RNA and Small RNA Isolation Kit (Biolabmix, Novosibirsk, Russia), according to the manufacturer’s protocol. Then, the purified tRNA was treated with 1U of RNase-free DNase I (1 U/μL) (Thermo Fisher Scientific, Waltham, MA, USA) in reaction buffer containing 40 mM Tris-HCl (pH 8.0), and 6 mM MgCl_2_. Reactions were incubated at 37 °С for 1 h. After extraction with phenol saturated with 0.1 M citrate buffer (pH 4.3) and isopropanol precipitation, tRNA was subjected to deacylation with 0.1 M Tris-HCl (pH 9.0) at 37 °С for 45 min. The pH was further decreased by the addition of 3 M NaOAc (pH 4.5) to its final concentration of 0.3 M. After isopropanol precipitation, uncharged tRNA was dissolved in nuclease-free water and dialyzed twice against 1000 volumes of ultrapure water.

### Rapid kinetics measurements

To monitor the time course of aminoacyl-tRNA interaction with the A-site of the ribosome, we rapidly mixed either (i) 0.1 µM initiation ribosome complexes containing BODIPY-Met-tRNA^fMet^ programmed with the MG mRNA with 1 µM Gly-tRNA^Gly^·EF-Tu·GTP ternary complex at 20 °C or (ii) 0.1 µM Gly-tRNA^Gly^(Prf16/17/20)·EF-Tu·GTP ternary complex with 0.4 µM initiation ribosome complexes programmed with the same MG mRNA at 20 °C. In both cases, samples were mixed in equal volumes (60 µL). Fluorescence was recorded using SX-20 stopped-flow spectrometer (Applied Photophysics, Leatherhead, UK). Proflavin fluorescence was excited at 460 nm, BODIPY FL fluorescence was excited at 470 nm. Fluorescence intensity was measured after passing a cut-off filter KV 495 nm (Schott, Mainz, Germany) in both cases. Time courses were obtained by averaging 5-7 individual traces. Data were evaluated by fitting to a single-exponential function with a characteristic apparent rate constant (*k_app_*), amplitude (A), and final signal amplitude (F_∞_) according to equation F = F_∞_ + A · exp(*k_app_* · t), where F is the fluorescence at time t. Where necessary, two exponential terms were used. All calculations were performed using the GraphPad Prism software (version 9.3.1, Dotmatics, Boston, MA, USA).

### Monitoring of EF-Tu coelution with the ribosomes

Reaction mixture containing 1 µM initiation complexes programmed with the MG mRNA and 2 µM Gly-tRNA^Gly^(Prf16/17/20)·EF-Tu·GTP ternary complex was incubated at 37 °C for 2 min followed by application to size exclusion chromatography on a BioSuite 450 Å HR SEC (8 μm, 7.8×300 mm) column (Waters, Milford, MA, USA) in buffer TAKM_7_. 1 pmol of the central fraction of 70S peak was separated by SDS-PAGE, and semidry transfer was performed for 30 min at 25 V using polyvinylidene difluoride membranes for western blotting (Bio-Rad Laboratories, Hercules, CA, USA). The membrane was blocked with 5% bovine serum albumin (BSA) and incubated with primary monoclonal anti-His_6_ antibodies (His-Tag Antibody, Affinity Biosciences, Shanghai, China) diluted 1:2500 in 5% BSA dissolved in TBS [20 mM Tris-HCl, 150 mM NaCl, pH 7.6] for 1 h at RT. After washes in TBST [20 mM Tris-HCl, 150 mM NaCl, 0.1% Tween-20, pH 7.6], the membrane was incubated with secondary horseradish peroxidase–conjugated antibodies (Goat anti-Mouse IgG (H+L) Secondary Antibody, HRP, Thermo Fisher Scientific, Waltham, MA, USA) diluted 1:5000 in 7% non-fat dry milk (Sigma-Aldrich, Burlington, MA, USA) dissolved in TBS for 1 h at RT. The bands were developed using the Clarity™ Western ECL Substrate (Bio-Rad Laboratories, Hercules, CA, USA) and visualized using the chemiluminescence mode in the ChemiDoc™ MP Imaging System (Bio-Rad Laboratories, Hercules, CA, USA).

## RESULTS

### Bottromycin A_2_ inhibits bacterial translation by targeting an unconventional site

The main aim of this study was to uncover the mechanism of action of the peptide antibiotic bottromycin A_2_ (BotA2, Figure 2A). Originally discovered in the 1950s, BotA2 has long been known to inhibit protein synthesis in bacterial cells [33]. However, the details of its action remained obscure. In addition, clinical application of BotA2 is limited due to the poor stability under physiological conditions, probably caused by hydrolysis of the methyl ester group [19]. Therefore, we were also interested to compare BotA2 and its demethylated counterpart – bottromycin A_2_ carboxylic acid (BotCA, Figure 2A) – in a cell-free bacterial *in vitro* translation system. By using the commercially available PURExpress system reconstituted from individual components purified from *Escherichia coli* cells, we observed a dose-dependent inhibition of protein synthesis by both BotA2 and BotCA, with the latter one being two times less efficient than the former one (IC_50_ = 6.4 ± 0.8 μM and 13.4 ± 2.0 μM, respectively, Figure 2B). At the same time, BotCA was previously reported to exhibit 64 times less potency against Gram-positive strains compared to BotA2 [19], suggesting that the main reason for the low *in vivo* activity of BotCA may be its impaired ability to penetrate bacterial cells rather than inability to suppress translation.

**Figure 2.**
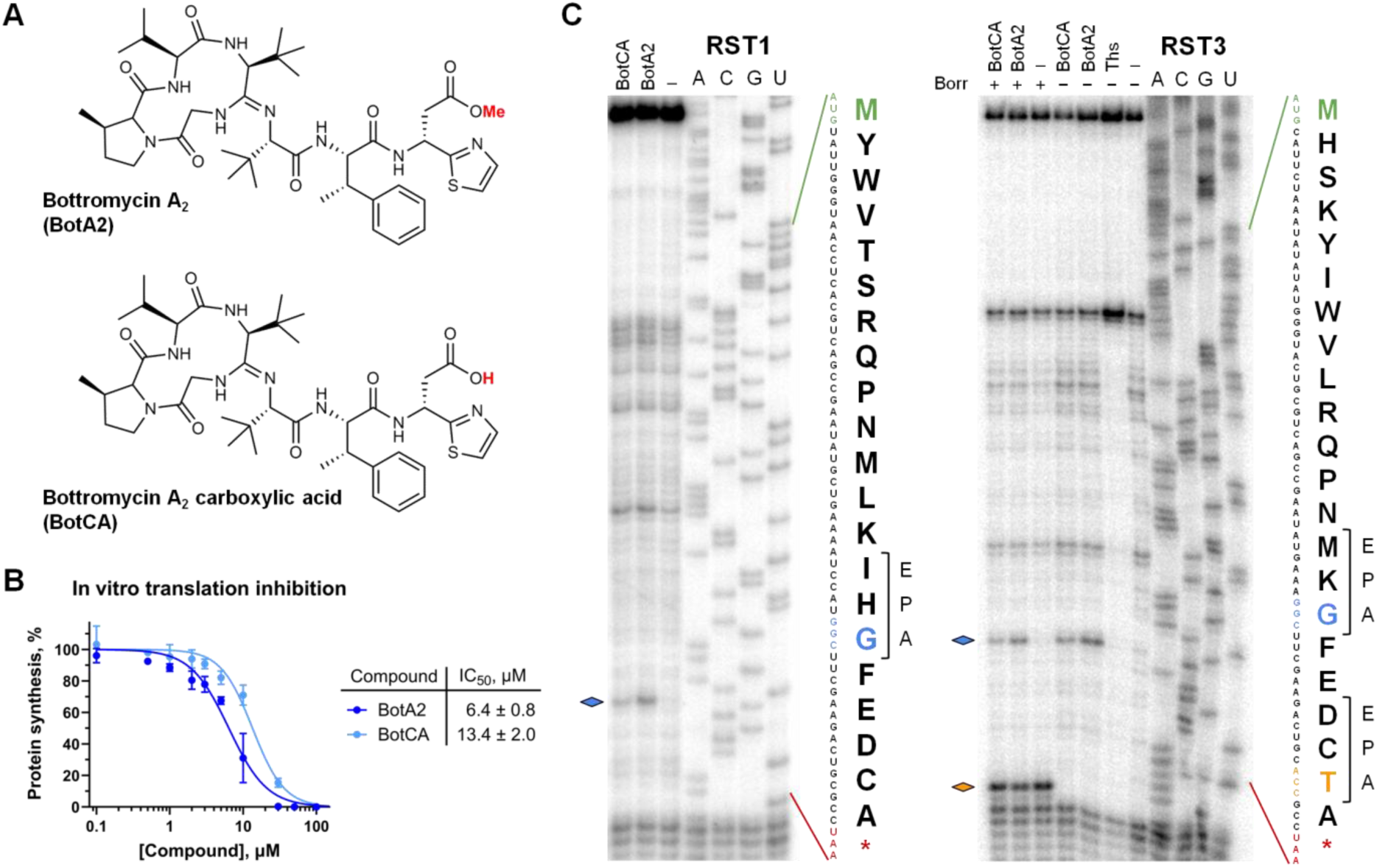
Bottromycin A_2_ inhibits bacterial translation in a context-specific manner. (**A**) Chemical structures of bottromycin A_2_ (BotA2) and its hydrolyzed counterpart referred to here as bottromycin A_2_ carboxylic acid (BotCA). (**B**) BotA2 and BotCA inhibit protein synthesis *in vitro*. The relative maximum Fluc accumulation rates are shown. Error bars represent standard deviation of the mean of at least two independent experiments. The calculated IC_50_ values and 95% confidence intervals are shown in the table. (**C**) Toe-printing analysis of BotA2 and BotCA on RST1 and RST3 mRNAs. Sequences of the corresponding ORFs and the encoded amino acids are shown on the right. Asterisk (*) in the translated sequence indicates a stop codon. Rhombuses point to the toe-printing bands corresponding to ribosomes stalled during translation. Codons located in the A-site of the stalled ribosomes are highlighted in the same color. Ths, thiostrepton, was used to indicate the translation start site. Borr, borrelidin. All antibiotics were tested at a final concentration of 50 μM.

By using another cell-free *in vitro* system based on the *E. coli* S30 extract, we observed a strong translation inhibition by BotA2 with IC_50_ = 1.0 ± 0.1 μM (Supplementary Figure S3), which is comparable to other translation inhibitors, such as tetracenomycin X [44], chloramphenicol [70], erythromycin [71], linezolid [72], and madumycin II [73] that display IC_50_ values of 1.5 μM, 2.1 μM, 0.32 μM, 1.1 μM, and 0.3 μM, respectively. In addition, we have tested if BotA2 and BotCA could interfere with eukaryotic mRNA translation. Both of them were found not to inhibit translation *in vitro* in HEK293T whole cell lysate (Supplementary Figure S4), meaning that bottromycin A_2_ specifically targets the machinery of bacterial translation.

Since BotA2 was previously assumed to interact with the 50S ribosomal subunit [34], we decided to test whether its binding site overlaps with that of other known ribosome-targeting antibiotics. By using agar diffusion assay, we screened our homemade collection of resistant mutants for the growth inhibition induced by BotA2. Surprisingly, we haven’t found any *E. coli* mutant resistant to BotA2 (Supplementary Figure S5), which is consistent with previous works showing no cross-resistance between BotA2 and erythromycin, as well as tetracycline [5,74]. Our data reveal that the BotA2 action is insensitive to the following nucleotide substitutions in the 23S rRNA ‒ U1782C, G2057A, A2058G, A2059G, A2062G, U2586G, U2586C, U2609G, while they confer strong resistance to erythromycin, chloramphenicol, or tetracenomycin X (Supplementary Figure S5).

In addition, we checked whether the Cfr-mediated methylation of A2503 in the 23S rRNA would confer resistance to BotA2, as it has been shown for chloramphenicol, lincosamides, oxazolidinones, pleuromutilins, and streptogramins A [75]. Antibacterial activity of BotA2 was measured against the *Bacillus subtilis* 168 strain engineered to express the *cfr* gene [52], compared to the susceptible strain (Supplementary Table S4). As expected, *cfr* expression resulted in an 8-fold increase in the minimum inhibitory concentration (MIC) of chloramphenicol (100 μM). At the same time, we observed no significant differences in MIC values for BotA2 between the two *B. subtilis* strains. This result implies that BotA2 does not interact with the A2503 residue. Our reasoning is additionally supported by one of the previous works, which revealed that BotA2 does not compete with chloramphenicol for the binding site [35]. Altogether, these findings suggest that BotA2, being a potent translation inhibitor, does not share binding site with classical 50S-targeting antibiotics, but rather possesses a unique binding site.

### Bottromycin A_2_ inhibits translation in a context-specific manner

Since the previous investigation reported that BotA2 inhibits aminoacyl-tRNA (aa-tRNA) binding in the A-site of ribosomes [32], we decided to visualize which step of translation indeed is inhibited by BotA2 using the toe-printing assay. Application of RST1 and RST3 mRNA templates revealed that BotA2 induces ribosome stalling during the elongation step, and specifically when a glycine (Gly) codon in the mRNA reaches the A-site of the ribosome (Figure 2C). This observation contradicts what we expected to see from a conventional inhibitor of aa-tRNA accommodation, like tetracycline, kirromycin, or lincosamides (lincomycin, clindamycin) [76]. They rather cause multiple toe-printing bands at the beginning of mRNA coding sequence, as in the case of 30S-targeting tetracycline, along with pronounced ribosomal arrest at the start codon [46]. Translation reactions on RST3 mRNA supplemented with borrelidine (Borr), which inhibits Thr-tRNA synthetase (Thr-RS), revealed that a fraction of ribosomes passes through the BotA2 stalling site and becomes trapped at the downstream site with Thr codon placed in the A-site (Figure 2C). We assume that BotA2 may reversibly bind to its target, as was previously suggested [35].

The obtained results prompt us to comprehensively examine the BotA2 context specificity. A recently developed high-throughput technique named toe-seq was applied for this purpose [60]. Toe-seq uses a library of short DNA templates containing a 30-nt randomized region within its ORF. *In vitro* expression of this library using a coupled transcription-translation system in the presence of BotA2, followed by treatment with AMV reverse transcriptase resulted in the generation of multiple parallel toe-prints. The corresponding cDNA fragments were subsequently subjected to next generation sequencing (NGS) in order to identify ribosome stalling sites and correlate them with the mRNA sequence. Toe-seq analysis was performed with 50 μM BotA2 in two biological replicates along with untreated control reactions. Datasets for antibiotic-treated samples were normalized to the controls and filtered according to a set of criteria (see Materials and Methods for details) to extract the most relevant data. The resulting *MaxStopProbability* scores, representing the relative efficiency of antibiotic-induced ribosome stalling, were well correlated between both replicates (Supplementary Figure S6). To determine whether specific sequence signatures are associated with the sites of BotA2-mediated translation stalling, we applied pLogo analysis [61] to 12,245 sites identified in two biological replicates. By looking at amino acid residues associated with codons located in the E-, P-, and A-sites of arrested ribosomes, we observed a strong tendency for the prevalence of Gly as the incoming amino acid (Figure 3A), which is in agreement with our earlier observations from toe-printing experiments (Figure 2C). Additionally, a marginal, but detectable enrichment was observed for Ala codons placed in the A-site. At the same time, there was no clear dependence of ribosome stalling on the last or penultimate amino acid residue of the nascent polypeptide chain. Fixing Gly or Ala at the position +1 of the pLogo plot revealed some minor preferences for amino acid residues at positions –1 and 0 (Figure 3B; Supplementary Figure S7). However the significance of these preferences is questionable and requires further validation.

**Figure 3.**
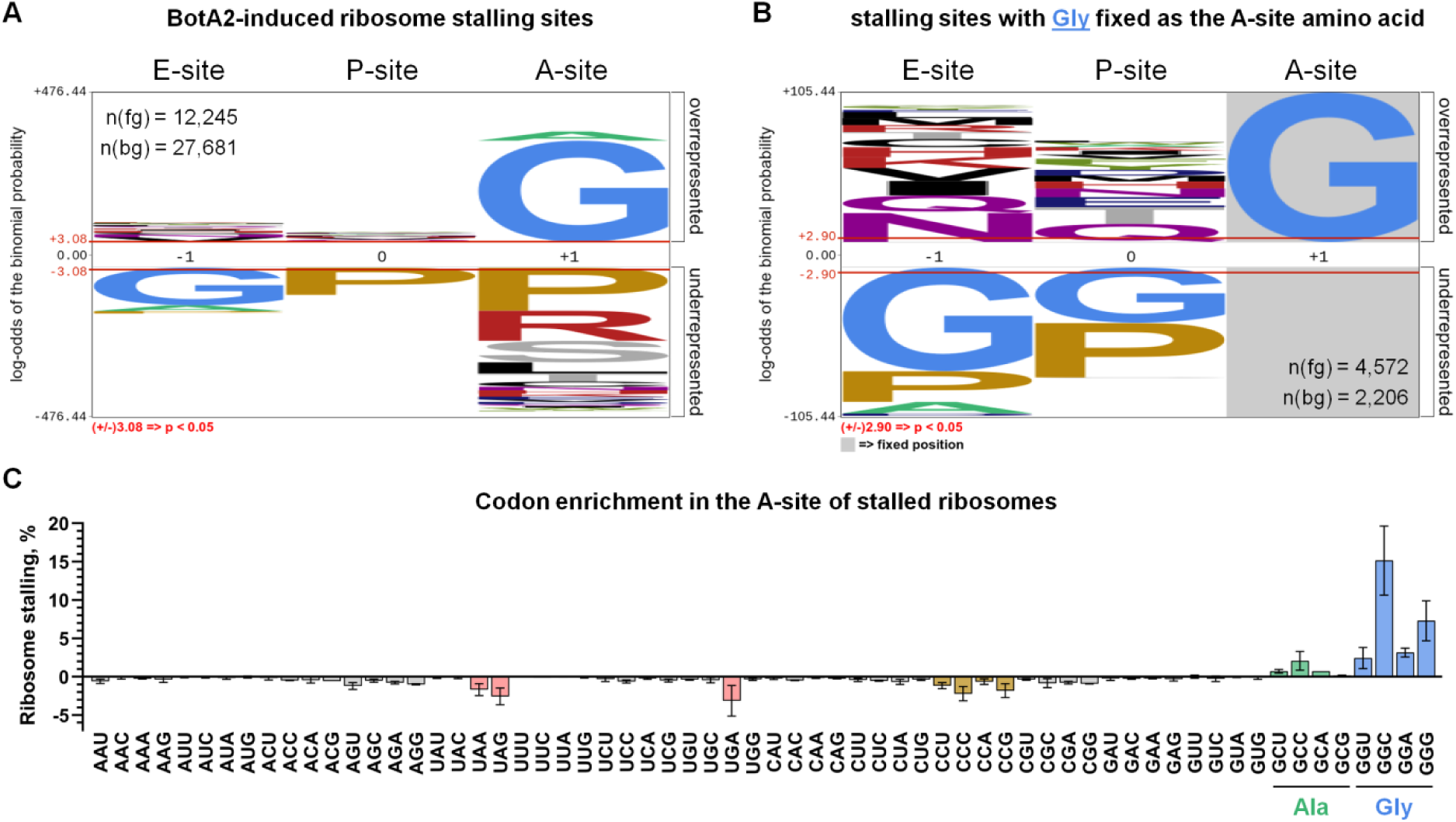
Toe-seq analysis of bottromycin A_2_ (BotA2) context specificity. (**A**) pLogo analysis of BotA2-induced ribosome stalling sites. Amino acids corresponding to codons located in the E-, P-, and A-sites of arrested ribosomes are indicated. The n(fg) and n(bg) values represent the number of foreground and background sequences used to generate the image, respectively. As the foreground, we used stalling sites identified in BotA2-treated samples. As the background, we used apparent stalling sites from samples that were not treated with antibiotics. The red horizontal bars on the pLogo correspond to p-value = 0.05. (**B**) pLogo analysis of a subset of BotA2-induced ribosome stalling sites, in which glycine (Gly, G) is fixed as the A-site amino acid (position +1). (**C**) Enrichment of codons occupying the A-site of the ribosomes stalled in the presence of BotA2. Enrichment is shown as the mean of normalized relative occurrence of codons among 12,463 identified stalling sites in two biological replicates. Codon frequency values for the untreated control samples were subtracted from the corresponding values for the BotA2-treated samples to account for possible codon bias. Therefore, the resulting negative values conform to underrepresented codons. Only codon positions 4-11 of the ORF are included in the analysis, as they correspond to the variable region of mRNA. Error bars indicate standard deviation.

We were also interested to analyze whether the efficiency of BotA2-induced ribosome stalling depends on individual codons more than on encoded amino acids. To this end, we calculated the relative occurrence of individual codons among the identified stalling sites and noticed that different A-site Gly codons exhibit different stalling frequencies, with GGC being the most efficient one, followed by GGG (Figure 3C). Similarly, among the A-site Ala codons, GCC was associated with more frequent translation arrest compared to GCU, GCA, and GCG. At the same time, we have not found any statistically significant enrichment of codons located in the E– and P-sites of stalled ribosomes. These data, along with the distribution of ribosome stalling sites across the toe-seq mRNA library, are provided in the Supplementary Excel Table. Altogether, the toe-seq analysis confirmed that BotA2 acts as a context-specific inhibitor of bacterial translation, arresting ribosomes specifically when a Gly codon occupies the A-site.

To validate observations made by means of toe-seq, we set up a conventional toe-printing system using a set of model mRNAs. Thus, short DNA templates encoding fMet-XXX-Gly peptides, were prepared to test if the identity of amino acid (or codon) located in the P-site is indeed irrelevant for BotA2-mediated ribosome stalling (Figure 4A). Since the GGC codon was associated with the most prominent stalling, we selected it to encode Gly. The XXX codons corresponding to the 2^nd^ amino acid in peptide were randomly selected so that some of them were associated with high *MaxStopProbability* scores, while others – with medium and low scores (Supplementary Figure S8). The toe-printing analysis of the generated DNA templates revealed that BotA2 causes ribosome stalling at the 2^nd^ position, almost regardless of the P-site codon. The absence of any reliable difference in BotA2-induced translation arrest on these templates is in agreement with our earlier toe-seq results (Figure 3).

**Figure 4.**
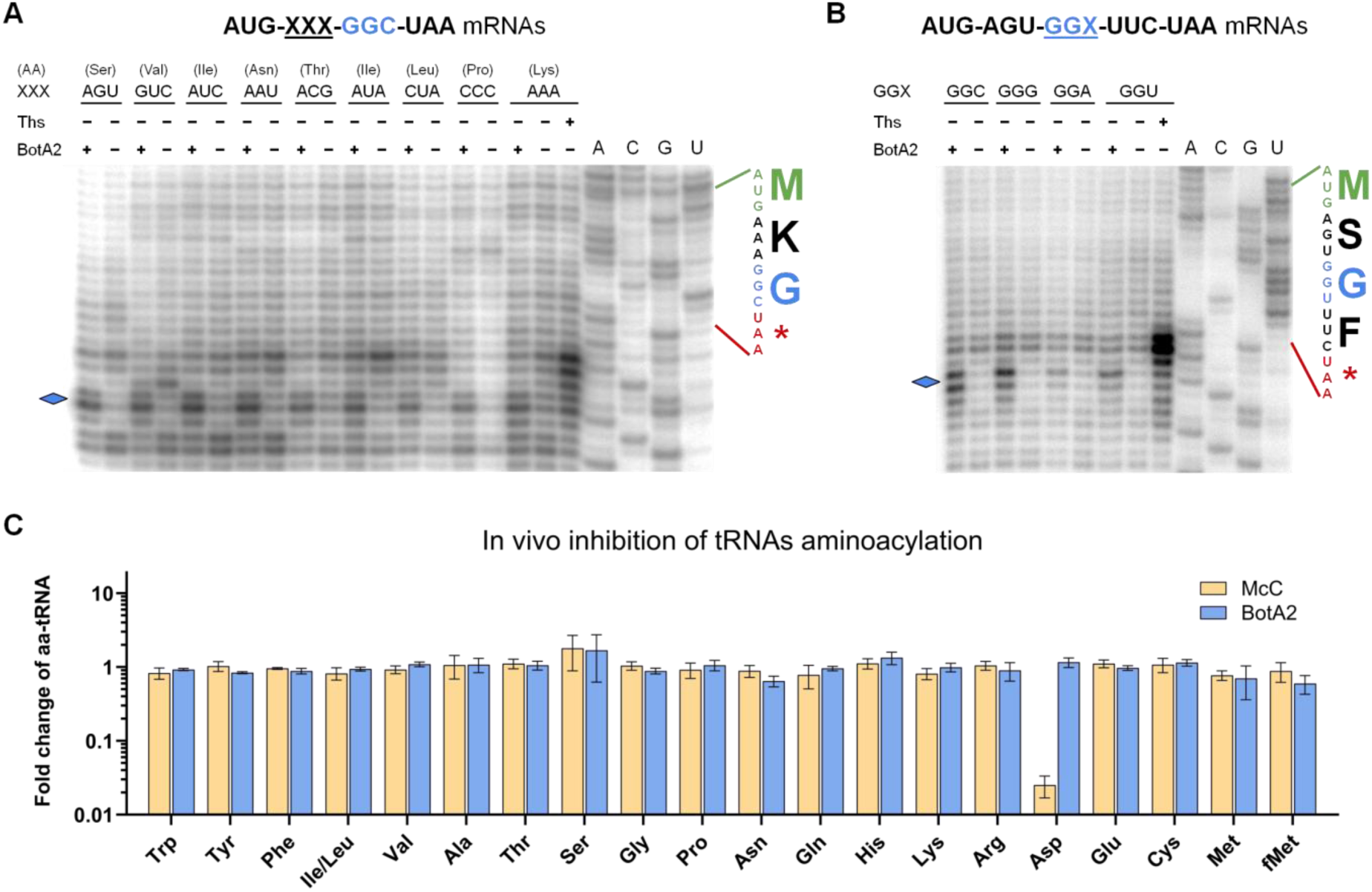
Context specificity of bottromycin A_2_ is unrelated to inhibition of tRNAs aminoacylation. (**A**) BotA2-induced ribosome stalling does not depend on the identity of the codon located in the P-site of the arrested ribosome. (**B**) BotA2-induced ribosome stalling depends on the identity of the Gly codon. (**A**,**B**) Toe-printing analysis of BotA2 on two sets of short mRNAs containing variable codons. Sequences of the corresponding ORFs and the encoded amino acids are shown on the right. Asterisk (*) indicates a stop codon. Rhombuses point to the toe-printing bands corresponding to BotA2-induced ribosome stalling. Ths, thiostrepton, was used to indicate the translation start site. All antibiotics were tested at a final concentration of 50 μM. (**C**) Addition of BotA2 does not change the amount of different aa-tRNAs in bacterial cells. Fold change in the level of aminoacyl-adenosines derived from aa-tRNAs that were purified from *E. coli lptD^mut^* cells treated with bottromycin A_2_ (BotA2) or microcin C (McC) is shown. The values obtained for antibiotic-treated samples were normalized to the values obtained from untreated cells. Error bars represent standard deviation of the mean of three independent experiments.

Next, we checked whether the efficiency of ribosome stalling induced by BotA2 depends on the identity of Gly codons, as predicted by toe-seq analysis (Figure 3C). Two sets of short DNA templates were prepared to be translated into fMet-Gly-Phe and fMet-Ser-Gly-Phe peptides, where Gly amino acid is encoded by different nucleotide triplets – GGC, GGG, GGA, or GGU. Both sets of templates showed that GGC and GGG codons were almost equally efficient, while GGU exhibited the least pronounced ribosomal arrest (Figure 4B; Supplementary Figure S9). Interestingly, that GGA demonstrated a strong toe-printing band, being placed at the 2^nd^ position in ORF (Supplementary Figure S9), while at the 3^rd^ position (Figure 4B) it was much less efficient at stalling ribosomes. This observation might be related to previous results showing that GGA considerably decreases translation efficiency in *E. coli* cells, when it is placed right after the start codon, and does not alter translation being placed at the downstream positions [77,78]. Probably, other factors unrelated to BotA2 enhance ribosome stalling at the start codon, when glycyl-tRNA is used as a substrate [77,79].

### Bottromycin A_2_ does not inhibit Gly-tRNA synthetase

One possible explanation of such a unique sequence specificity of BotA2 action could be if BotA2 inhibits aminoacylation of tRNA^Gly^. To test this hypothesis, we employed two different approaches. Typical inhibitors of aminoacyl-tRNA synthetases are known to induce RelA-dependent synthesis of guanosine tetraphosphate (ppGpp) (e.g., protein kinase HipA [80], microcin C [81], mupirocin [82], and serine hydroxamate [83]). Obviously, amino acid starvation also leads to a shortage of free ternary complexes along with the accumulation of deacylated tRNAs. The RelA protein was found to specifically recognize an uncharged tRNA in the A-site of a stalled ribosome, acting as a sensor of amino acid deficiency [84,85]. Upon activation, RelA converts cellular ATP and GTP/GDP to the alarmone ppGpp [86], thus inducing the stringent response aimed at upregulation of genes involved in amino acid biosynthesis, protein hydrolysis, and transition to a so-called “hibernation” state as a means of energy conservation and survival [87–89].

By using the metabolic labeling of BotA2-susceptible *E. coli lptD^mut^* strain (Supplementary Table S4), we examined whether BotA2 would lead to the accumulation of ppGpp. Bacterial cells were grown in the medium supplemented with [^32^P]-radiolabeled orthophosphoric acid, followed by treatment with BotA2. As positive controls, we used microcin C (McC) and mupirocin (Mup) known to inhibit Asp-tRNA synthetase [90] and Ile-tRNA synthetase [91], respectively. After 30 min incubation, the labeled nucleotides were extracted with formic acid and separated via thin-layer chromatography (TLC). As shown in Supplementary Figure S10, only McC and Mup, but not BotA2, provoked the accumulation of ppGpp in cells. This observation suggests that BotA2 does not increase the concentration of deacylated tRNAs and hence does not induce the stringent response.

In order to directly assess if BotA2 affects the intracellular pool of aa-tRNAs, global analysis of tRNA aminoacylation level (GATRAL) [63] was applied to *E. coli lptD^mut^* treated with BotA2. Treatment with microcin C (McC) was used as a positive control. Total tRNA was isolated after 30 min incubation with antibiotics, followed by *N*-acetylation of aa-tRNAs, required to stabilize amino acid residues bound to 3’-terminal adenosines. Then, the samples were digested with a mixture of RNase I and RNase T1 to obtain *N*-acetylated aminoacyl-adenosines (Ac-aa-Ade) that can be quantitatively analysed by LC-MS. The calculated fold change in the amount of each Ac-aa-Ade, relative to the untreated cells, reflects the abundance of the corresponding aa-tRNA in the cells (Figure 4C). As expected, McC resulted in a substantial decrease in the level of Asp-tRNA^Asp^ (see Asp in Figure 4C), while BotA2 showed no differences even with respect to Gly-tRNA^Gly^ (see Gly in Figure 4C). This analysis indicates that BotA2 does not affect aminoacylation of tRNA^Gly^, as well as other tRNAs.

Beyond its potential to inhibit Gly-tRNA synthetase, BotA2 could, in theory, directly target aminoacylated tRNA, analogous to GNAT-family toxins that specifically modify the aminoacyl moiety [92]. However, direct and specific interaction between this small antibiotic molecule and the acceptor stem of aa-tRNA seems mechanistically impossible. In addition, any non-*N*-acetylating modifications of amino acid residues of aa-tRNA would be expected to perturb the levels of acetylated glycyl-adenosines (Ac-Gly-Ade) used in the GATRAL assays, while the structural features of BotA2 (see Figure 2A) preclude its functioning as an acetylating agent.

Altogether, these findings strongly suggest that BotA2 neither inhibits Gly-tRNA synthetase, nor directly modifies Gly-tRNA^Gly^.

### Bottromycin A_2_ does not cause mistranslation of Gly codons

Another possible explanation could be if BotA2 specifically prevents Gly-tRNA^Gly^ from binding to the A-site of the ribosome, making it excessively available for erroneous binding of near-cognate aa-tRNAs. In this case, BotA2 should provoke translational misreading of Gly codons similarly to other error-inducing antibiotics [93]. To assess the ability of BotA2 to decrease the fidelity of translation, we exploited a set of reporter constructs encoding β-galactosidase with different substitutions for the catalytic residue Glu537. Upon normal translation, reporter cells produce the defective protein, which does not catalyze the hydrolysis of the X-Gal substrate. Although treatment with certain antibiotics (e.g., kanamycin or streptomycin), which increase the translational error rate, allows Glu to be “mistakenly” incorporated at the 537^th^ position resulting in the active β-galactosidase, which can be detected by an insoluble, indigo blue colored product [49]. By using agar diffusion assay, we tested BotA2, BotCA, and reference antibiotics on these reporter bacteria (Supplementary Figure S11). As expected, we observed blue halos upon treatment with kanamycin and streptomycin, while rifampicin, known to inhibit bacterial RNA polymerase [94], showed no coloration. Likewise, BotA2 and BotCA exerted no blue halos, even when tested on pJC27-GGA and pJC27-GGG reporters, in which the 537^th^ GAA codon was replaced by GGA or GGG, respectively. This observation led us to conclude that BotA2 does not provoke the misreading of Gly codons.

### Bottromycin A_2_ does not interfere with the delivery of Gly-tRNA^Gly^ to the A-site of the ribosome

Looking for a particular step of translation inhibited by BotA2, we next decided to check if it can specifically interfere with the delivery of Gly-tRNA^Gly^ to the A-site of the ribosome. For this purpose, we applied a modified toe-printing assay, where translation reactions were additionally treated with the RelE toxin, known to cleave mRNA codons in the ribosomal A-site (Figure 5A) [95]. Since the hydrolytic activity of RelE requires a vacant A-site, RelE-printing represents a useful tool to monitor the occupancy of the A-site during translation [59]. Two mRNAs, whose translation is known to be highly susceptible to BotA2-induced stalling, were selected for RelE-printing analysis of BotA2 action (Figure 5B). As expected, we observed the RelE-prints originating from initiating and terminating ribosomes (see cleavage at the 2^nd^ and 5^th^ codons, Figure 5B), since 70S initiation complex, pre-termination and post-termination complexes are characterized by available A-sites [96]. Noteworthy that the reaction supplemented with thiostrepton (Ths) exerted only one RelE-print related to ribosomes trapped at the start codon, which is consistent with the antibiotic’s mode of action [97,98]. At the same time, no RelE-prints corresponding to mRNA cleavage at Gly codons were observed. These data suggest that BotA2 allows delivery of Gly-tRNA^Gly^ to the A-site of the ribosome, thereby preventing the association of RelE and subsequent mRNA cleavage at Gly codons.

**Figure 5.**
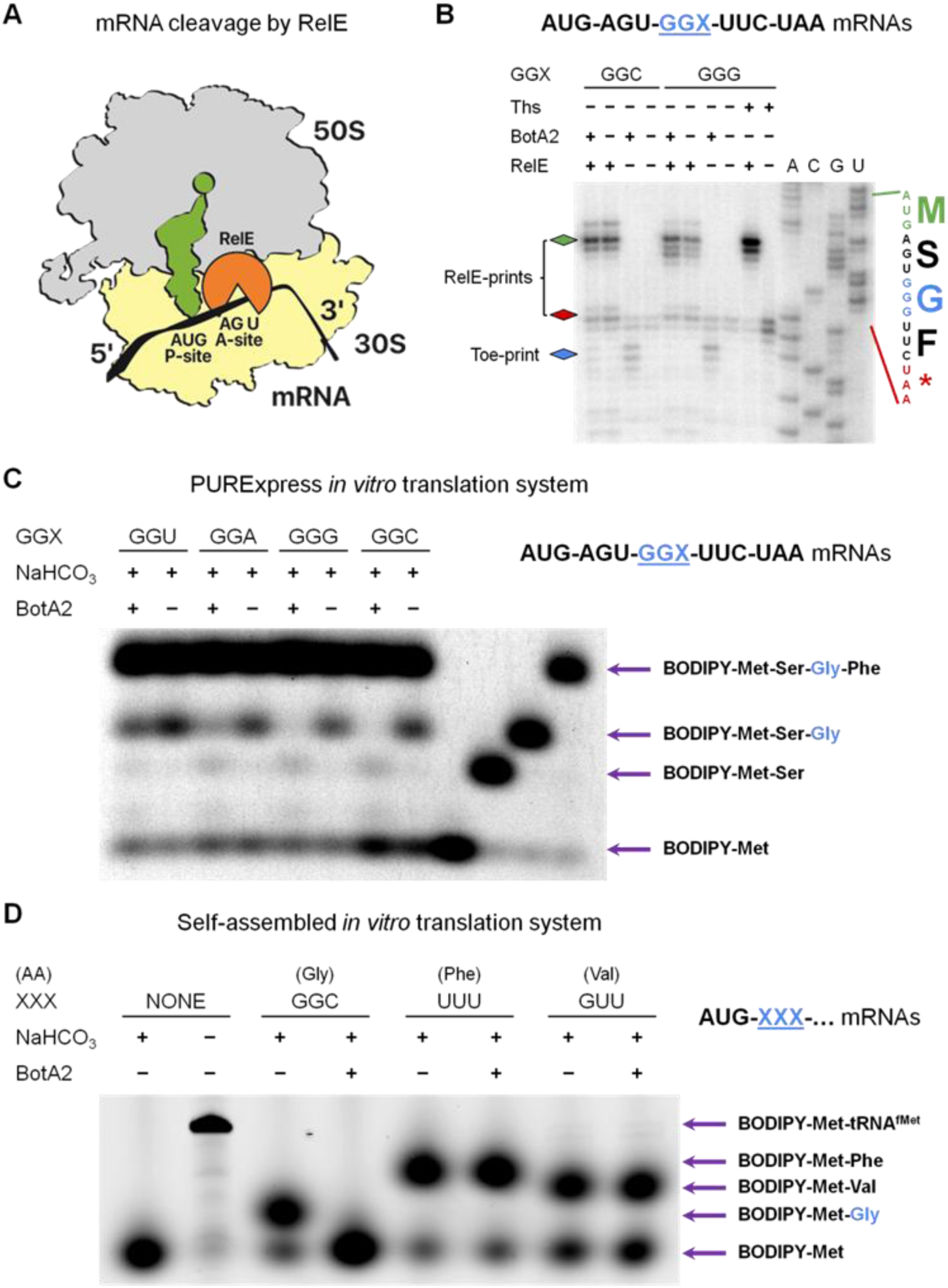
Bottromycin A_2_ allows delivery of Gly-tRNA^Gly^ to the A-site of the ribosome, but interferes with the peptide-bond formation. (**A**) Schematic representation of RelE-mediated cleavage of mRNA. RelE binds in the ribosomal A-site and cleaves mRNA after the 2^nd^ or 3^rd^ nucleotide in the A-site codon [95]. RelE does not cleave free mRNA, as well as when the A-site is occupied with aa-tRNA. (**B**) RelE-printing analysis of BotA2 action on two short mRNAs encoding fMet-Ser-Gly-Phe peptides. Toe-printing reactions supplemented with the RelE toxin indicate the position of ribosomes with the vacant A-site. Sequences of the corresponding ORFs and the encoded amino acids are shown on the right. Asterisk (*) in the translated sequence indicates a stop codon. Blue rhombus points to the toe-printing bands corresponding to BotA2-induced ribosome stalling. Green and red rhombuses point to the RelE-printing bands corresponding to mRNA fragments cleaved at the 2^nd^ and 5^th^ codons. Ths, thiostrepton, was used to indicate the translation start site. All antibiotics were tested at a final concentration of 50 μM. (**C**,**D**) *In vitro* synthesis of fluorescently labeled peptides depending on the presence of BotA2 using either PURExpress (**C**) or self-assembled (**D**) translation system. In (**C**), each reaction was supplied with BODIPY-labeled fMet-tRNA^fMet^ as a sole source of *N*-formylmethionine, purified uncharged tRNAs, and amino acids ‒ Ser and Gly. Phe was not added intentionally to “freeze” ribosomes at the Gly codon. Note that due to the Phe impurities contained in the kit, the full-length peptide was synthesized regardless. BotA2 was tested at a final concentration of 50 μM. In (**D**), BODIPY-Met-Gly/Phe/Val dipeptides were formed upon addition of a pre-assembled cognate ternary complex to the 70S initiation complexes programmed with MG (AUG-GGC), MF (AUG-UUU), or MV (AUG-GUU) mRNA and containing BODIPY-Met-tRNA^fMet^ in the P-site. Where indicated, ternary complexes were additionally incubated with BotA2, which was tested at a final concentration of 100 μM. After incubation, translation reactions were treated with NaHCO_3_ to hydrolyze peptidyl-tRNA and release synthesized peptides.

### Bottromycin A_2_ inhibits peptidyl transfer to Gly-tRNA^Gly^

Having determined that the delivery of Gly-tRNA^Gly^ was not suppressed by BotA2, we sought to uncover which of the following steps ‒ Gly-tRNA^Gly^ accommodation, peptide-bond formation, or translocation ‒ is directly affected by BotA2. Some of the preceding works argued in favour of inhibition of translocation [36,39], while others stated that inhibition of peptide-bond formation may be the primary action of BotA2 [37,38]. To discriminate between these two main hypotheses, we decided to monitor peptide formation in the presence of BotA2. A recently developed method based on *in vitro* synthesis of fluorescently labeled peptides [66,69] was applied for this purpose.

Short mRNAs from the previous experiments were used to express BODIPY-labeled peptides upon treatment with BotA2 (Figure 5C). To reduce the effect of multiple-round translation, we decided to “freeze” translating ribosomes at the Gly codon by lacking Phe ‒ the last amino acid in the tetrapeptide. To our surprise, the full-length BODIPY-Met-Ser-Gly-Phe tetrapeptide was synthesized anyway, probably due to the Phe impurities contained in the PURExpress Δ(aa, tRNA) Kit. Some amino acids or aminoacyl-adenylates (aa-AMP) may co-purify with aminoacyl-tRNA synthetases [99] during the kit preparation, thereby compromising an experiment using short mRNA templates. Nevertheless, once all Phe impurities were completely used up, we were able to observe the formation of truncated peptides originating from arrested ribosomes. If BotA2 inhibited translocation, then we would expect to see the formation of the BODIPY-Met-Ser-Gly tripeptide to the same extent regardless of the presence of BotA2. In reality, we observed the inhibition of BODIPY-Met-Ser-Gly synthesis in all reactions supplemented with BotA2, regardless of the identity of Gly codon (Figure 5C). In agreement with our earlier results (Figure 4B), GGG and GGC codons were associated with the highest inhibitory effect, while GGU was the least efficient. Along with the inhibition of BODIPY-Met-Ser-Gly synthesis we also observed the accumulation of the BODIPY-Met-Ser dipeptide, indicating that BotA2 primarily inhibits peptide-bond formation rather than translocation.

However, due to impurities in the PURExpress *in vitro* translation system, the degree of inhibition of peptide transfer to Gly-tRNA^Gly^ was difficult to assess. Thus, we decided to additionally validate our findings using a reconstituted *in vitro* translation system consisting of individual purified components [66]. The reactions were assembled by combining initiating 70S ribosomes attached to the mRNA, bearing the AUG start codon in the P-site followed by the codon directing incorporation of Gly (GGC), Phe (UUU), or Val (GUU) amino acid, and the cognate ternary complex (aa-tRNA·EF-Tu·GTP), bearing Gly-tRNA^Gly^, Phe-tRNA^Phe^, or Val-tRNA^Val^, respectively (Figure 5D). Consistent with the previous experiment, we observed that BotA2 was able to completely prevent the formation of BODIPY-Met-Gly, whereas it had no effect on the synthesis of BODIPY-Met-Phe and BODIPY-Met-Val.

Altogether, our findings strongly suggest that BotA2 specifically impairs incorporation of glycine into a growing peptide chain.

### Bottromycin A_2_ prevents accommodation of Gly-tRNA^Gly^ in the peptidyl transferase center

Since peptidyl transfer to Gly-tRNA^Gly^ located in the A-site is impaired in the presence of BotA2, we were thinking whether BotA2 directly inhibits peptide-bond formation or if it actually prevents the accommodation of Gly-tRNA^Gly^ in the peptidyl transferase center (PTC) prior to transpeptidation. Therefore, we set up to visualize the approaching of acceptor stem of Gly-tRNA^Gly^ to the PTC. To this end, we used a stopped-flow fluorescence detection assay to monitor the delivery of glycine in the form of the ternary complex Gly-tRNA^Gly^·EF-Tu·GTP into the vicinity of BODIPY-labeled methionine residue of initiator tRNA located in the P-site of the ribosome (Figure 6A). Interestingly enough, for BotA2-treated complexes we did not observe the conventional increase of fluorescent signal, characteristic for the A-site accommodation of aa-tRNA acceptor end [65]. This observation allowed us to suppose that BotA2 does not abolish peptidyl transferase reaction *per se*, rather impairs the preceding step of amino acid delivery to the PTC.

**Figure 6.**
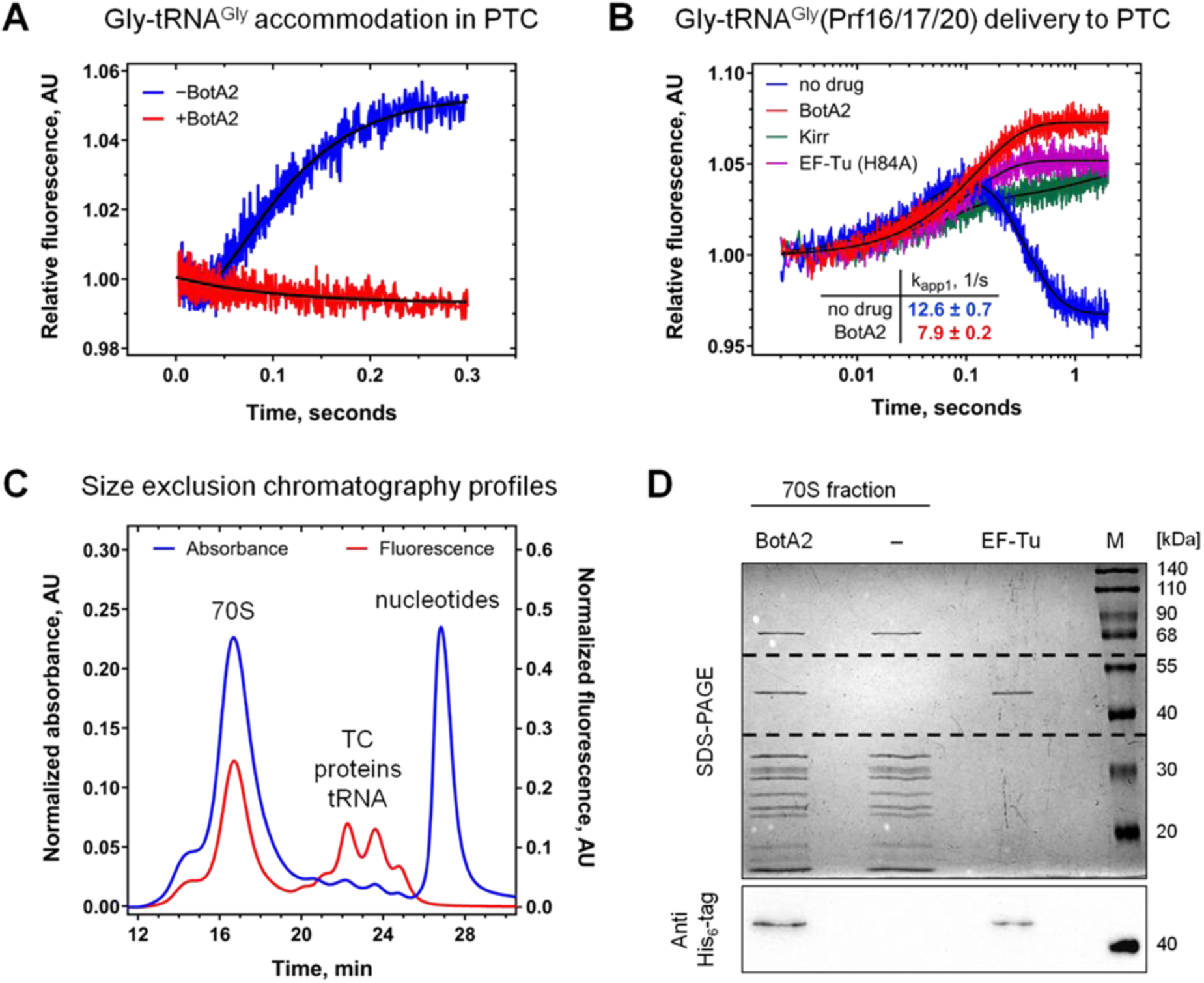
Bottromycin A_2_ prevents accommodation of Gly-tRNA^Gly^ in the peptidyl transferase center (PTC) by trapping ternary complex on the ribosome. (**A**,**B**) Stopped-flow experiments showing that BotA2 prevents accommodation of Gly-tRNA^Gly^ in the ribosomal A-site. (**A**) Pre-steady state kinetics of Gly-tRNA^Gly^ accommodation monitored by fluorescence change of BODIPY-Met-tRNA^fMet^ in the P-site of the 70S initiation complex (0.1 μM) upon mixing with the ternary complex Gly-tRNA^Gly^·EF-Tu·GTP (1 μM) in the absence (blue) or presence of BotA2 (red). (**B**) Pre-steady state kinetics of Gly-tRNA^Gly^ binding to the A-site monitored by fluorescence change of the ternary complex Gly-tRNA^Gly^(Prf16/17/20)·EF-Tu·GTP (0.1 μM) upon interaction with the 70S initiation complex (0.4 μM) containing fMet-tRNA^fMet^ in the P-site. Time courses were obtained in the absence (blue) or presence of BotA2 (red), as well as in the presence of kirromycin (Kirr, green) or modified variant of EF-Tu (His84Ala) deficient on GTP hydrolysis (purple). (**C**,**D**) BotA2 impairs EF-Tu dissociation from the translating ribosome. (**C**) Profiles of size exclusion chromatography of the 70S initiation complex after incubation with the ternary complex Gly-tRNA^Gly^(Prf16/17/20)·EF-Tu·GTP in the presence of BotA2. Absorbance values were measured at 260 nm (blue), with the scale displayed on the left Y-axis. Fluorescence was measured using excitation at 460 nm and detection at 510 nm (red), with the scale displayed on the right Y-axis. TC, ternary complexes. (**D**, upper panel) SDS-PAGE of 1 pmol of His-tagged EF-Tu (lane EF-Tu) and 1 pmol of 70S peak fraction corresponding to ribosomal complexes assembled in the absence (lane ‒) or presence of BotA2 (lane BotA2). (**D**, lower panel) Immunoblotting of the area between the dashed lines with anti-His_6_ antibodies. Where indicated, BotA2 was tested at a final concentration of 100 μM.

To monitor the effect of BotA2 on the delivery of glycine residue in real time, we continued to use a stopped-flow technique and prepared a modified tRNA^Gly^, labeled with the fluorescent dye at the elbow region ‒ tRNA^Gly^(Prf16/17/20), as interaction of proflavin-labeled tRNA with the ribosomal A-site is well characterized and can be divided into several substeps [100]. An increase in a typical biphasic fluorescence change cumulatively represents the initial binding of the ternary complex to the ribosome and subsequent codon recognition, which induces GTPase activation and results in GTP hydrolysis by EF-Tu. The successive decay of the high fluorescence intermediate is related to the release of aa-tRNA from the GDP-bound form of EF-Tu and accommodation of the aa-tRNA in the A-site of the ribosome (Figure 6B). Exponential fitting of the fluorescence curves allowed us to measure the rate constants of reactions preceding GTP hydrolysis and subsequent tRNA accommodation. Remarkably, BotA2 slightly decreased the rate of ternary complex binding (*k_app1_*(no drug) = 12.6 ± 0.7 s^−1^ versus *k_app1_*(BotA2) = 7.9 ± 0.2 s^−1^), and completely abolished the following step – accommodation of Gly-tRNA^Gly^.

The obtained results imply two major possibilities. The first one is that BotA2 hinders productive tRNA accommodation by forming a steric obstacle or changing the conformation of the ribosome that closes the accommodation corridor for tRNA. However, it is difficult to imagine that the effect is pronounced in the case of the smallest amino acid, glycine, whereas bulky amino acids like phenylalanine or valine effectively pass through and participate in peptidyl transferase reaction. The second scenario would be tRNA retention in a bent conformation in the A/T state with anticodon bound to mRNA codon and the acceptor stem interacting with EF-Tu, which leads to the arrest of the ternary complex on the ribosome. To discriminate between these two possibilities, we performed size exclusion chromatography of arrested ribosomal complexes, which allowed us to separate all the components according to their hydrodynamic radii. As expected, 70S ribosomes eluted first showing the largest peak visualized by absorbance at 260 nm (blue profile, Figure 6C).

Proflavin-labeled Gly-tRNA^Gly^, detected by fluorescence, was present both in the ribosomal fraction and in the central part of the chromatogram corresponding to ternary complexes (TC), proteins and tRNAs. To check whether EF-Tu was trapped on the ribosome upon addition of BotA2, we analyzed the ribosomal fraction by SDS-PAGE and immunoblotting with anti-His_6_ antibodies against His-tagged EF-Tu (Figure 6D). Clear EF-Tu band appeared only in the case of antibiotic treated ribosomes, suggesting that BotA2 impairs EF-Tu dissociation from the translating ribosome.

Ternary complex retention on the ribosome was commonly associated with EF-Tu inability either to hydrolyze GTP or to undergo subsequent conformational rearrangements.

Previously, that was observed in the case of mutated variant of EF-Tu (His84Ala) deficient on GTP hydrolysis [101,102] or experiments with antibiotic kirromycin that traps EF-Tu in an active GTP-bound conformation even after GTP hydrolysis, preventing tRNA from accommodation [101,103] (see comparison in Figure 6B). Notably, kirromycin stalls various ternary complexes regardless of tRNA identity, which is in agreement with the antibiotic binding site at the interface of domain I (also known as G-domain) and domain III of EF-Tu, owing to which kirromycin interferes with conformation transition of the factor [104,105]. If the mechanism of BotA2 action were associated with violation of EF-Tu-mediated GTP hydrolysis or subsequent factor rearrangements, then most likely BotA2 would not have specificity for one particular amino acid. Preference of BotA2 for glycine could be rationalized by another, previously undescribed, option with potential binding of the antibiotic in the proximity of amino acid binding pocket of EF-Tu, located in domain II at a distance from the GTP binding site. As EF-Tu is evolutionary designed to accept 21 distinctive in size and charge amino acids, one could anticipate simultaneous binding of the smallest amino acid and BotA2, whereas all other amino acids won’t left enough space for the antibiotic. The proposed immobilization of the glycine residue in the presence of BotA2 would definitely prevent tRNA from its full accommodation in the ribosomal A-site.

Structurally the situation is reminiscent of interaction of glycine and another antibiotic chloramphenicol leading, however, to the opposite effect. Chloramphenicol, being a canonical peptidyl transferase antibiotic, fails to inhibit transpeptidation when the incoming A-site substrate is glycine, because fully accommodated glycine residue can co-exist in the A-site with the ribosome-bound chloramphenicol [106], whereas larger residues either adopt improper orientation or displace the antibiotic [107]. Thus, we suppose, that BotA2 exhibits specificity to glycine due to the ability of the antibiotic to bind simultaneously with the smallest amino acid to EF-Tu. Being bound, BotA2 impairs EF-Tu dissociation from the translating ribosome, resulting in the observed inhibition of Gly-tRNA^Gly^ accommodation in the PTC.

## DISCUSSION

In this study, we have examined the mode of action of bottromycin A_2_ (BotA2) ‒ a peptide antibiotic originally discovered in the 1950s as a potent inhibitor of translation machinery. For a long time, the details of its mechanism of action remained obscure with a number of conflicting reports being published. Here, we showed that BotA2 has a unique context specificity with regard to the mRNA coding sequence. We revealed that BotA2 induces stalling of elongating ribosomes exclusively when a Gly codon enters the A-site of the ribosome, with stalling efficiency independent of codons located in the P– and E-sites. Looking for possible explanations of such a unique context specificity, we have proved that BotA2 does not interfere with aminoacylation of tRNAs^Gly^ and does not provoke the misreading of Gly codons. At the same time, the formation of the peptide bond between the incoming Gly-tRNA^Gly^ and the peptidyl group of the P-site bound peptidyl-tRNA in the presence of BotA2 was completely abolished. Furthermore, we demonstrated that BotA2 traps Gly-containing ternary complexes on the ribosome with the tRNA anticodon bound to the mRNA in the A-site and unaccommodated acceptor stem of Gly-tRNA^Gly^, most likely retained in complex with EF-Tu.

In addition, we showed that the binding site of BotA2 differs from that of other known ribosome-targeting antibiotics. Even though one of the previous works [34] suggested that BotA2 binds 50S ribosomal subunit, it does not exhibit cross-resistance to erythromycin, chloramphenicol, tetracenomycin X, lincosamides and other antibiotics targeting peptidyl transferase center (Supplementary Figure S5; Supplementary Table S4). Selection of resistant mutants is considered the gold standard when it comes to identification of antibiotic’s binding site. We tried several times to select *E. coli* mutants resistant to BotA2, and we always received either a bacterial lawn or no clones (data not shown). While these observations highlight therapeutic potential of BotA2, our findings suggest that BotA2 binds in the unique site, different from that of classical 50S-targeting translation inhibitors.

In-depth structural analysis is required to shed light on how BotA2 interacts with its target, whereas genetic approaches are essential to validate this target. While preparing the manuscript, we figured out that another group of researchers was working with BotA2 and tried to obtain the structure of it bound to ribosomal complexes [108]. They showed for the first time that BotA2 directly binds ribosome associated ternary complexes at the interface of domain I and domain II of EF-Tu and thus prevents EF-Tu dissociation from the ribosome.

Aminoacyl-tRNA in such a complex adopts a bent conformation characteristic for the A/T state, which is consistent with the inhibition of Gly-tRNA^Gly^ accommodation in the peptidyl transferase center, that we observed (Figure 6). As anticipated, they proved that BotA2 can bind only ternary complexes bearing a Gly-tRNA^Gly^ or at least Ala-tRNA^Ala^, while other aa-tRNAs would form a steric clash with the antibiotic. These pioneering structural findings, additionally supported with genetic experiments confirming EF-Tu being a direct target of BotA2, complemented our observations and inspired us to verify that BotA2 traps Gly-containing ternary complexes on the ribosome in our self-assembled *in vitro* translation system. Thus, the structural and genetic work of our colleagues in combination with our biochemical and biophysical results allows one to conclude that bottromycin A2 possesses a novel, previously undescribed mechanism of action based on the context-specific immobilization of ternary complexes on elongating ribosomes.

Searching for an explanation, why BotA2 exhibits some preferences towards different Gly codons, we analyzed if it might be related to codon usage or the abundance of corresponding tRNAs (Supplementary Table S5). In *E. coli* cells codons GGU and GGC are overrepresented among highly expressed genes, while codons GGA and GGG are extremely underrepresented [109]. This correlates well with intracellular concentrations of cognate tRNAs^Gly^. In total, three isoaccepting tRNAs^Gly^ are encoded in the *E. coli* genome [110].

Codons GGC and GGU are recognized by tRNA^Gly^_GCC_ (referred to here as Gly3), GGA is recognized exclusively by tRNA^Gly^_UCC_ (referred to here as Gly2), while GGG is decoded by both tRNA^Gly^_UCC_ and tRNA^Gly^_CCC_ (referred to here as Gly1) (Supplementary Table S5). In agreement with codon usage, Gly3 is the most abundant tRNA^Gly^ in *E. coli* cells compared to Gly1 and Gly2. Probably, GGA and GGG codons were not accidentally depleted during evolution, since they resemble the Shine-Dalgarno sequence and thus may hinder productive translation elongation [111]. However, this does not explain the observed preferences of BotA2.

The only logical explanation seems to be that the strength of the codon-anticodon interaction determines the efficiency of BotA2-induced ribosome stalling (Supplementary Figure S12). Probably, the perfect codon-anticodon Watson-Crick pairing stabilizes a Gly-tRNA^Gly^ in the arrested complex. For example, Gly1 and Gly3 perfectly pair to GGC and GGG codons, respectively, through the formation of 9 hydrogen bonds in total. Meanwhile, Gly2 forms only 8 hydrogen bonds upon pairing with GGA or GGG. However, the 5-methylaminomethyl-modification of uridine (mnm^5^U) at the first anticodon position of Gly2 was previously shown to additionally stabilize pairing to both A and G [49,112]. Hence, the GGA and GGG recognition by Gly2 leads to the formation of slightly less stable codon-anticodon duplexes. In contrast, GGU recognition by Gly3 contributes to the formation of the least stable (weak) complex maintained by 8 hydrogen bonds with the G•U wobble base pair (Supplementary Figure S12). Altogether, Gly codons can be arranged in the following order, depending on the strength of codon-anticodon interactions: GGC ≥ GGG > GGA > GGU. In general, this order correlates well with BotA2 preferences and indirectly explains why GGU is the most frequent Gly codon in highly expressed genes (∼60%). One may assume that a less stable base pairing would increase the probability of Gly-tRNA^Gly^ dissociation from the ribosome in the form of a complex with EF-Tu and BotA2. Presumably, inhibition of aa-tRNA accommodation and subsequent peptide-bond formation compels ternary complexes bearing less “sticky” Gly-tRNAs^Gly^ to undergo repeated rounds of association/dissociation, thus increasing the ribosome’s chances to accidentally bypass the stalling motif. The similar mechanism was recently shown for chloramphenicol [113]. This would explain why the GGU codon was the least efficient one, while ribosomal complexes with GGC and GGG codons were the most stable ones upon BotA2-induced stalling.

Special attention should be paid to the fact that the *btmD* gene, that encodes the bottromycin precursor peptide, starts from the ATG-GGA-CCC-… evolutionary conserved codons, further converted into the fMet-Gly-Pro-… amino acid sequence [9]. Since BotA2 specifically traps ribosomes decoding a Gly codon, we assume that BotA2 might limit its own production via a negative feedback loop. As predicted earlier, the bottromycin (*btm*) gene cluster contains only one self-resistance gene *btmA*, which encodes a major facilitator superfamily (MFS) transporter, probably responsible for BotA2 secretion. It’s quite likely that when the concentration of BotA2 in the fermentation broth reaches a certain level, the intracellular concentration of the antibiotic also starts to increase until equilibrium. High levels of BotA2 inside the producing cells may specifically lead to downregulation of *btmD* translation, and, as a result, to inhibition of further BotA2 production for the sake of survival. Moreover, the BtmH epimerase was shown to bind excessive BotA2 [16], which might provide an additional protection against self-poisoning [8]. On the one hand, BtmH can titrate the excess BotA2 with high affinity, thus lowering its intracellular concentration. While, on the other hand, this BotA2 directly inhibits BtmH and thereby prevents further maturation of new BotA2 molecules [16]. Overall, these mechanisms might in fact control BotA2 yield upon fermentation with the aim of avoiding self-harm.

As previously shown, BotA2 is not stable upon physiological conditions and degrades in plasma faster than it reaches the focus of inflammation [18]. At the same time, a range of BotA2 derivatives were synthesized with some of them demonstrating increased stability and promising antibacterial properties [19]. We hope that our work will facilitate new attempts of obtaining improved derivatives of BotA2, and new BotA2-based medicines would be available for patients suffering from multidrug resistant pathogens in the foreseeable future.

## ACKNOWLEDGEMENTS

We express our sincere gratitude to Prof. Rolf Müller and Dr. Joy Birkelbach at the Helmholtz Institute for Pharmaceutical Research Saarland (HIPS) and Helmholtz Centre for Infection Research (HZI) (Saarbrücken, Germany) for providing pure bottromycin A_2_, as well as bottromycin A_2_ carbocylic acid. We thank Prof. Daniel N. Wilson (Institute for Biochemistry and Molecular Biology, University of Hamburg, Hamburg, Germany) for cooperation in early stages of the project development and valuable discussions. We are especially grateful to Dr. Ilya A. Osterman, who originally conceived this project, for supporting early aspects of this work. We should also acknowledge the impact of the work of Yury S. Polikanov (University of Illinois, Chicago, IL, USA) and his colleagues, who have shared their structural and genetic data, demonstrating the role of EF-Tu in BotA2 mechanism of action, ahead of publication. We also thank Andrey G. Tereshchenkov and Dr. Natalia V. Sumbatyan at the Lomonosov Moscow State University (Moscow, Russia) for helpful suggestions and attempts to reveal the binding site of bottromycin A_2_. We thank Elizaveta A. Razumova for assistance with *E. coli ΔtolC* pJC27 reporter cells. We thank Dr. Dmitry E. Andreev at the Belozersky Institute of Physico-Chemical Biology (Lomonosov Moscow State University, Moscow, Russia) for providing the purified RelE protein and discussing results. We thank Tinashe P. Maviza for providing *E. coli* SQ171 strains transformed with different pLK35 and pAM552 plasmids. We thank the staff at the Skoltech Advanced Mass Spectrometry Core Facility (Moscow, Russia) for performing LC-QTOF-MS analysis, especially Maria Zavialova and Dmitriy Maslov. Preparation of uncharged total tRNA, metabolic labeling, and analysis of aminoacylation levels of tRNAs were performed by Alexei Livenskyi, whose work was supported by the grant of Russian Science Foundation [24-14-00181].

## AUTHOR CONTRIBUTIONS

Inna A. Volynkina: Validation, Formal analysis, Data curation, Software, Investigation (*in vitro* translation assays, toe-printing, RelE-printing, misreading reporter assay, synthesis of BODIPY-peptides with PURExpress system), Writing—original draft, Visualization, Project administration. Aleksandr A. Grachev: Formal analysis, Investigation (synthesis of BODIPY-peptides with self-assembled *in vitro* translation system, rapid kinetics measurements, monitoring of EF-Tu coelution with the ribosomes). Alexei Livenskyi: Methodology, Validation, Formal analysis, Data curation, Investigation (metabolic labeling assay, GATRAL), Writing—original draft, Visualization. Daria K. Yagoda: Investigation (*in vitro* translation assays, toe-printing, RelE-printing, growth inhibition of resistant mutants, synthesis of BODIPY-peptides with PURExpress system, solid-phase extraction). Pavel S. Kasatsky: Investigation (tRNA^Gly^ purification, self-assembled *in vitro* translation system). Olga A. Tolicheva: Investigation (synthesis of BODIPY-peptides with self-assembled *in vitro* translation system). Ekaterina S. Komarova: Methodology, Investigation (toe-seq). Alexey E. Tupikin: Investigation (NGS). Vera A. Alferova: Methodology, Validation, Data curation, Resources, Investigation (RP-HPLC purification of BotA2 and BotCA), Writing—original draft, Visualization. Anastasiia O. Karakchieva: Investigation (toe-printing, MIC measurement, solid-phase extraction). Arina A. Nikandrova: Data curation, Investigation (solid-phase extraction, analytical RP-HPLC). Mikhail V. Biryukov, Yuliya V. Zakalyukina, and Lubov V. Dorofeeva: Methodology, Investigation (cultivation of the producing strain), Writing—review & editing. Yuriy A. Ikhalaynen and Igor A. Rodin: Validation, Investigation (LC-HRMS/MS). Dmitrii A. Lukianov: Methodology, Resources, Writing—review & editing, Project administration. Marsel R. Kabilov: Formal analysis, Resources, Data curation, Software. Alena Paleskava: Methodology, Validation, Formal analysis, Data curation, Writing—original draft, Visualization, Funding acquisition. Andrey L. Konevega: Conceptualization, Resources, Writing—review & editing, Supervision, Project administration. Petr V. Sergiev: Conceptualization, Formal analysis, Resources, Writing—review & editing, Supervision, Project administration. Olga A. Dontsova: Resources, Supervision, Funding acquisition.

## SUPPLEMENTARY DATA

Supplementary Data are available at NAR online.

## CONFLICT OF INTEREST

The authors declare no conflict of interest.

## FUNDING

This work was supported by Russian Science Foundation (RSF), grant number 21-64-00006-P to O.A.D (*in vitro* translation, toe-printing, RelE-printing, toe-seq, experiments with bacterial cells) and grant number 25-14-00253 to A.P. (experiments on self-assembled translation system, pre-steady state kinetics, and EF-Tu trapping). The funders had no role in study design, data collection and analysis, decision to publish, or manuscript preparation.

## DATA AVAILABILITY

The toe-seq sequencing data have been deposited to the NCBI BioProject database under accession number PRJNA1304769. Bioinformatics scripts are available at https://github.com/kabilov/Publication_scripts/tree/main/2024_Toe-seq. Other relevant data that support this study are available in the main manuscript, supplementary information, or from the corresponding authors upon reasonable request.

## Notes

https://www.ncbi.nlm.nih.gov/bioproject/PRJNA1304769

### Competing Interest Statement

The authors have declared no competing interest.

### Summary of Updates

This version of the manuscript has been revised to update the list of authors and information about purification of bottromycin A2 and its carboxylic acid from culture broth of the producing strain.

