## Supplementary Materials for "Insights into the bottromycin A_2_ mechanism of action"

**Figure S1.** RP-HPLC data for BotA2 and BotCA purification

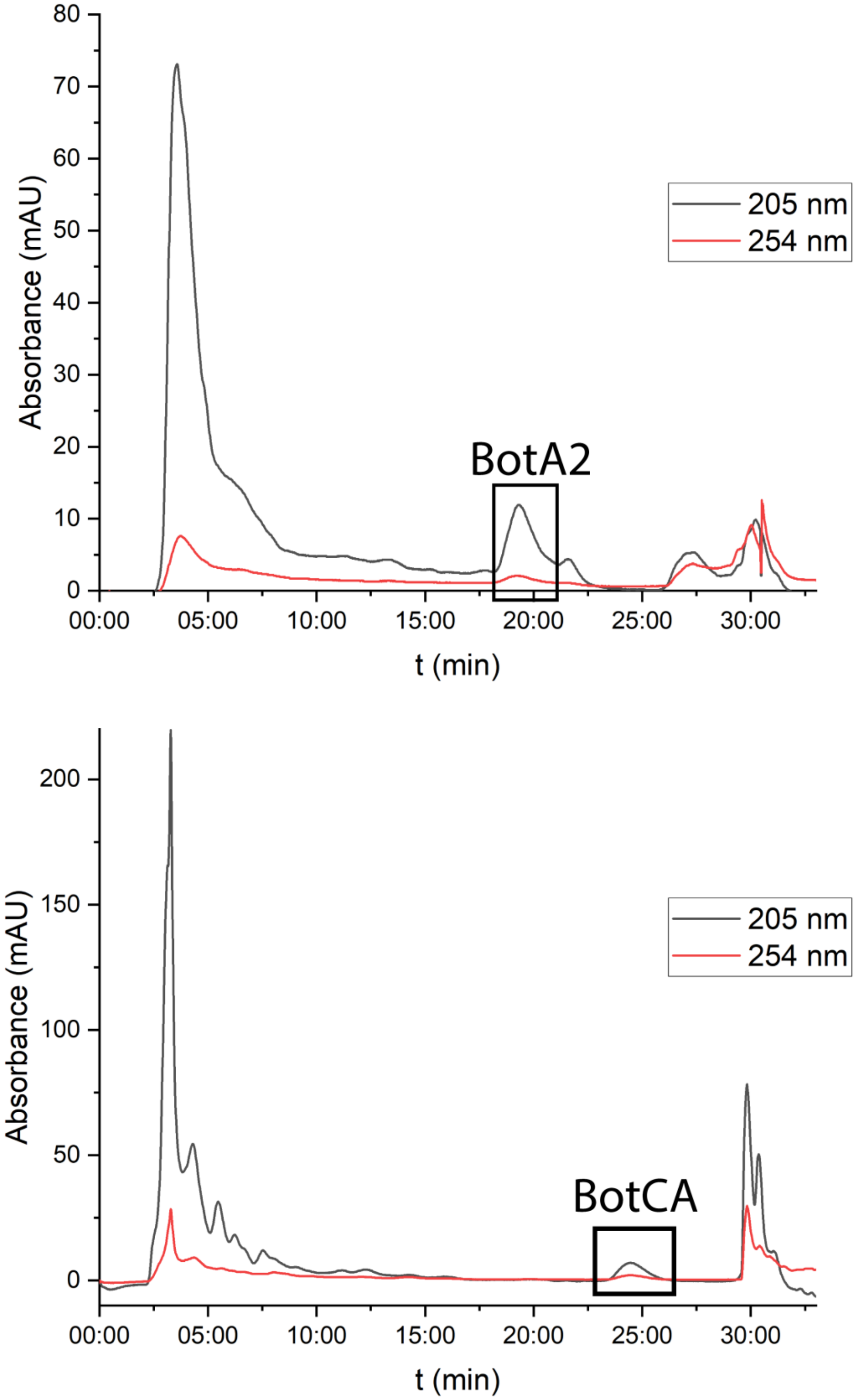

**Figure S2.** HRMS/MS fragmentation for BotA2 and BotCA

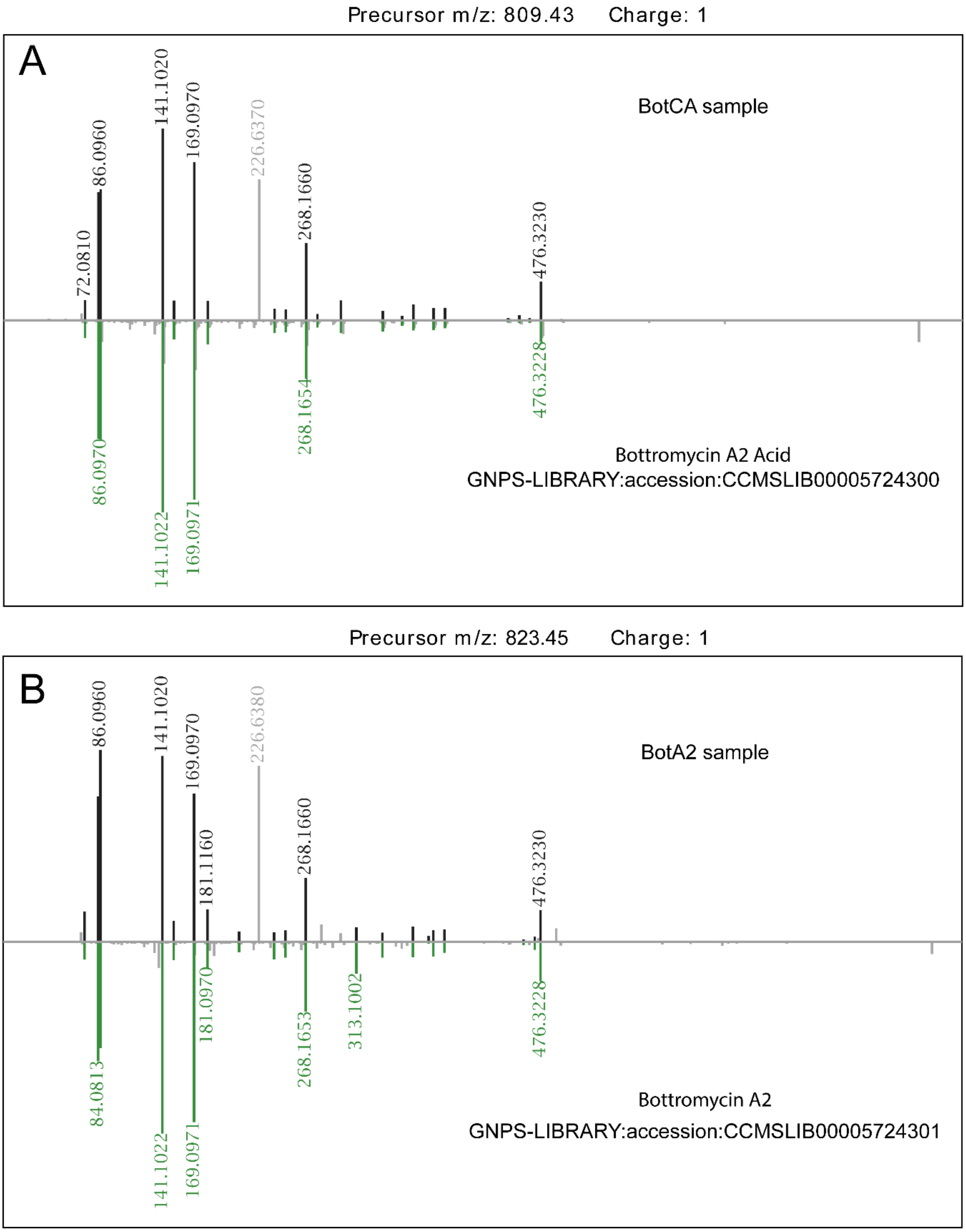

Comparison of the MS/MS fragmentation spectra of BotCA (**A**) and BotA2 (**B**), illustrated using the GNPS molecular networking platform [1]. The analysis was carried out for the $[M+H]^+$ adducts, with precursor masses indicated in the figure. The experimental fragmentation patterns were mirrored against reference spectra from the GNPS library: bottromycin $A_2$ acid (accession CCMSLIB00005724300) and bottromycin $A_2$ (accession CCMSLIB00005724301).

**Figure S3.** Inhibition of protein synthesis *in vitro* by BotA2 in a bacterial cell-free system

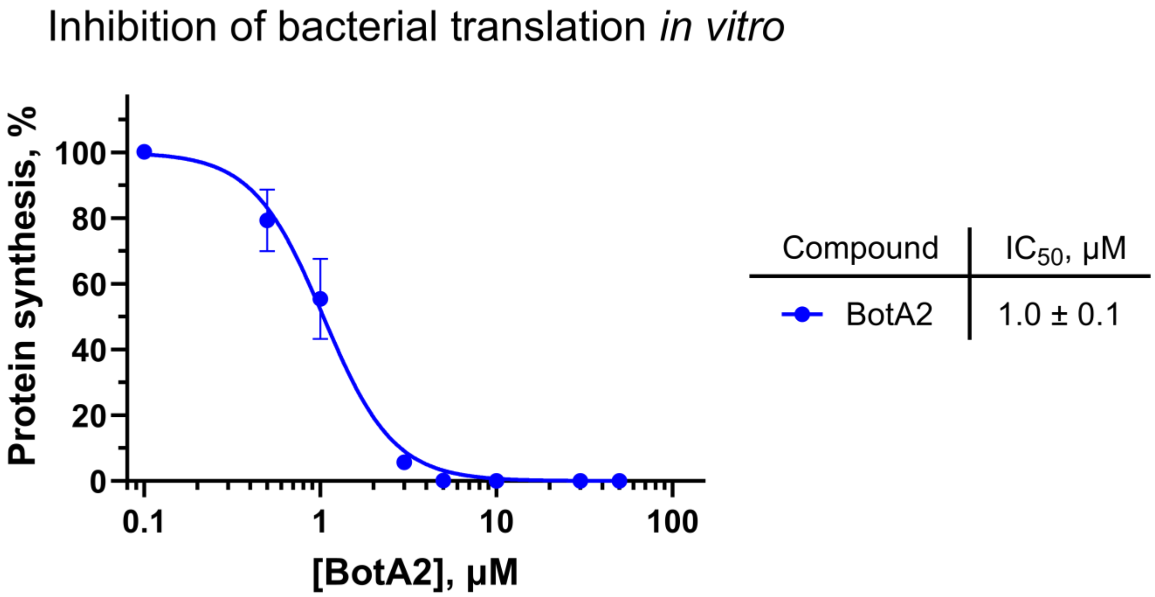

Inhibition of protein synthesis by increasing concentrations of bottromycin $A_2$ (BotA2) was asessed *in vitro* in a cell-free bacterial translation system using commercially available *E. coli* S30 extract. The relative maximum Fluc accumulation rates are shown. Error bars represent standard deviation. All reactions were repeated at least two times. The calculated $IC_{50}$ values and 95% confidence intervals are shown in the table.

**Figure S4.** Inhibition of protein synthesis *in vitro* by BotA2 and BotCA in HEK293T whole cell lysate system

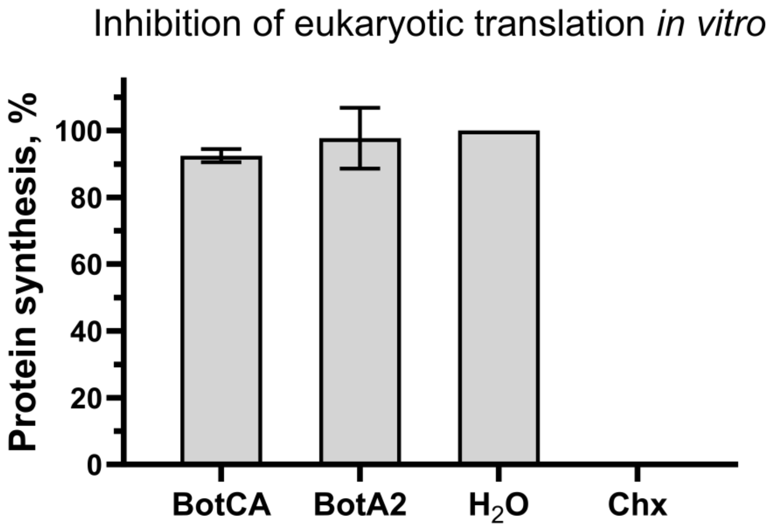

Inhibition of protein synthesis was assessed *in vitro* in a mammalian cell-free translation system based on HEK293T whole cell extracts. The relative maximum Fluc accumulation rates are shown. Chx, cycloheximide, was used as a positive control. BotA2, bottromycin $A_2$. BotCA, bottromycin $A_2$ carboxylic acid. Error bars represent standard deviation. All compounds were tested at a final concentration of 50 μM. All reactions were repeated at least two times.

**Figure S5.** Antibacterial activity of BotA2 against *Escherichia coli* cells carrying mutations in the 23S rRNA gene.

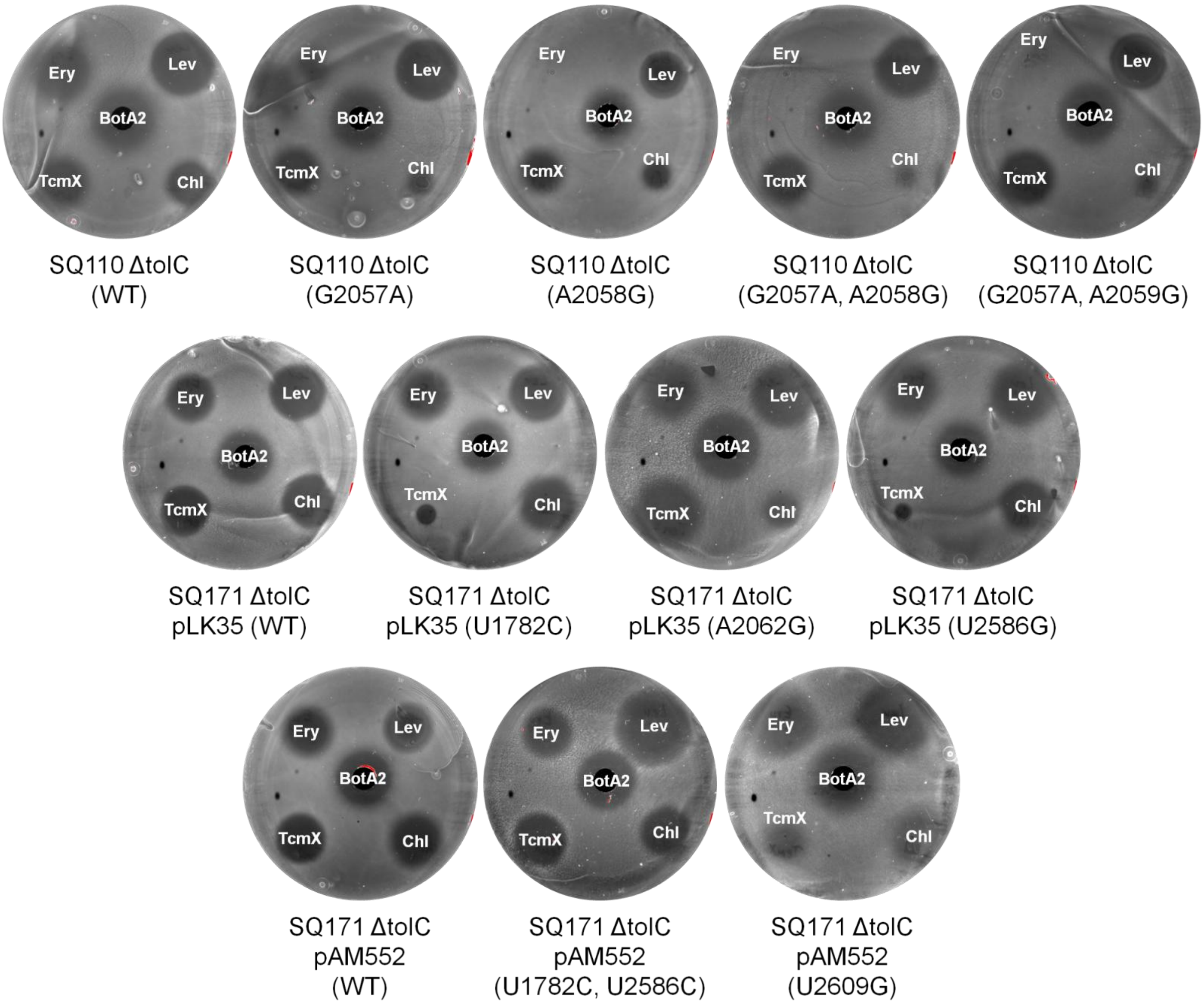

Agar diffusion assay reveals the activity of bottromycin $A_2$ (BotA2) against a panel of *E. coli* strains resistant to typical 50S-targeting antibiotics, such as erythromycin, tetracenomycin X, and chloramphenicol. Resistance is defined by point mutations in the 23S rRNA gene. The corresponding nucleotide substitutions are indicated in brackets. WT, wild-type 23S rRNA gene. The following antibiotics were applied on the surface of an agar plate coated with SQ110 ΔtolC or SQ171 ΔtolC cells: erythromycin (Ery, 5 µg), levofloxacin (Lev, 25 ng), tetracenomycin X (TcmX, 5 µg), chloramphenicol (Chl, 2 µg). In the center of each agar plate, a well was made and filled with 50 µL of 1 mM BotA2 dissolved in 30% acetonitrile aqueous.

**Figure S6.** Correlation analysis of *MaxStopProbability* scores between two independent toe-seq experiments.

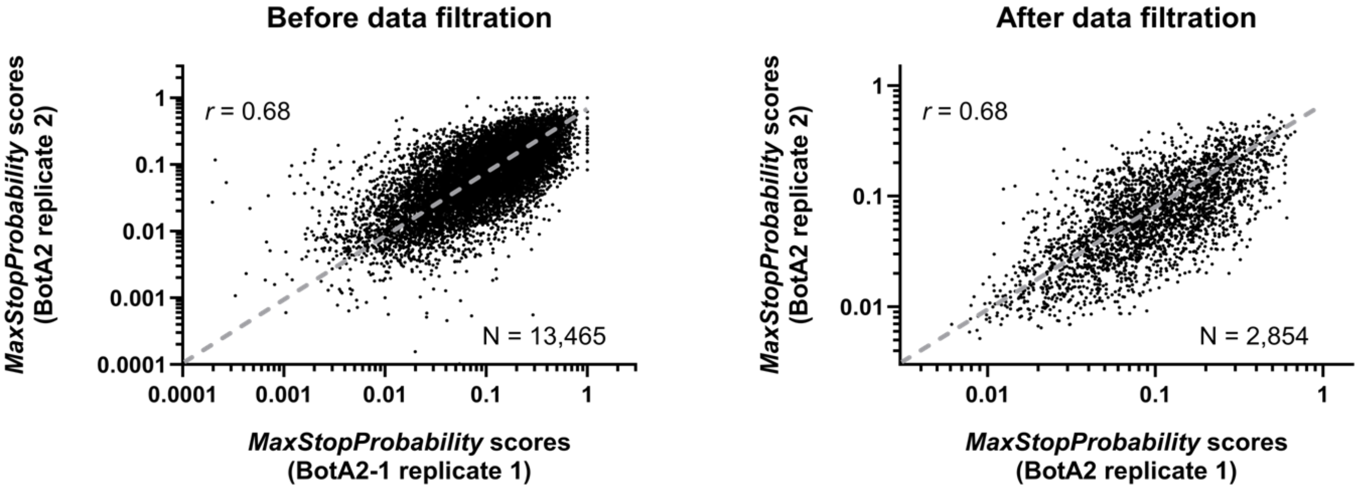

Toe-seq analysis of BotA2 was performed in two biological replicates. *MaxStopProbability* scores have been calculated to reveal antibiotic-specific ribosome stalling sites. A subset of mRNAs associated with coincident stalling sites (*StopPosition*) in two replicates was selected for correlation analysis. Only mRNAs with *MaxStopProbability* scores greater than 0 were considered. The number of such mRNAs (N) is presented on each plot. Spearman correlation coefficient (*r*) of the corresponding *MaxStopProbability* scores is presented on each plot, p-value < 0.0001 (two-tailed). The trend line is shown as a gray dashed line.

**Figure S7.** pLogo analysis of ribosome stalling sites with alanine fixed as the incoming A-site amino acid.

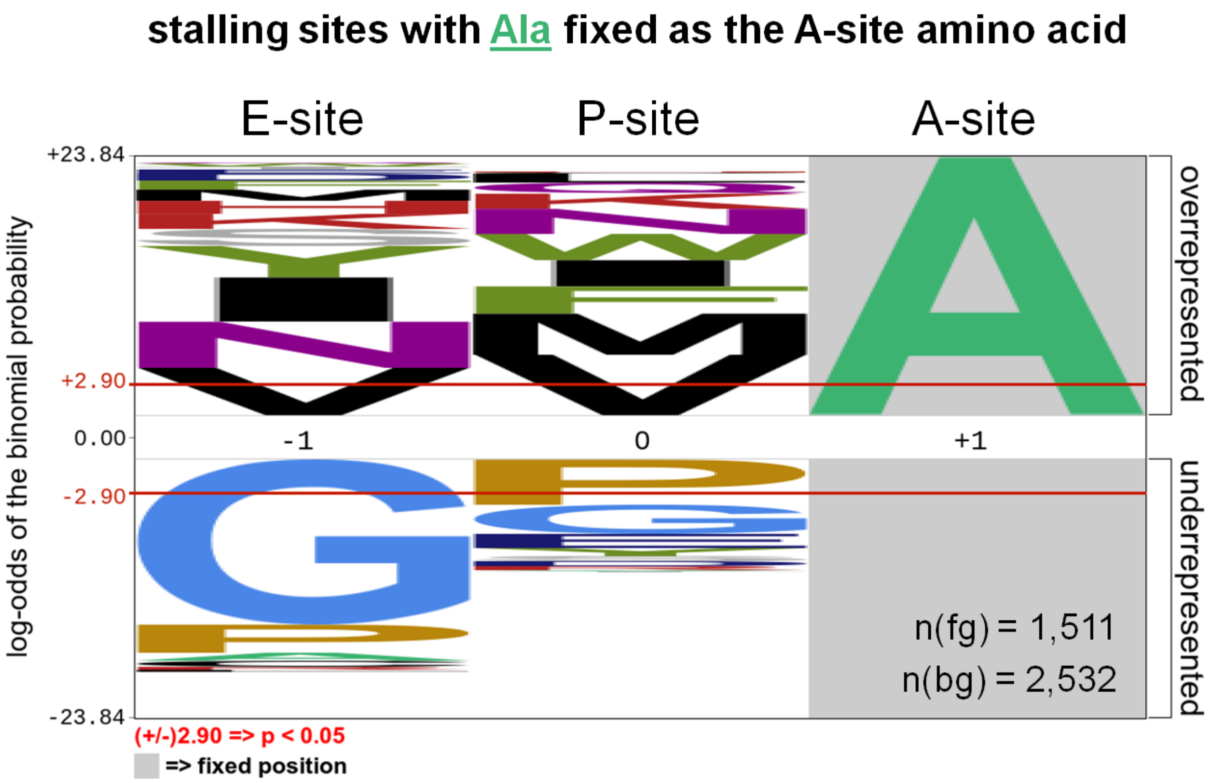

pLogo analysis of a subset of BotA2-induced ribosome stalling sites, in which alanine (Ala, A) is fixed as the incoming A-site amino acid (position +1). Amino acids corresponding to codons located in the E- and P-sites of arrested ribosomes are indicated. The n(fg) and n(bg) values represent the number of foreground and background sequences used to generate the image, respectively. The red horizontal bars on the pLogo correspond to p-value = 0.05.

**Figure S8.** *MaxStopProbability* scores of AUG-XXX-GGC ribosome stalling sites, depending on the identity of the XXX codon.

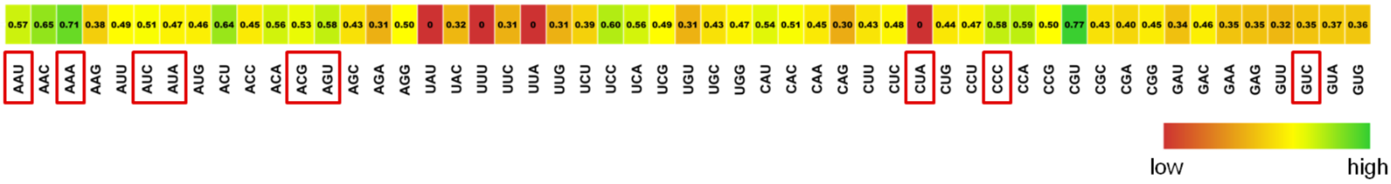

*MaxStopProbability* scores corresponding to different AUG-XXX-GGC ribosome stalling sites, where XXX is the second codon of the open reading frame (ORF) and does not encode Gly or Ala, are indicated within the cells of the heatmap. The values were calculated as the maximum among both toe-seq datasets derived from BotA2-treated samples. A *MaxStopProbability* score of 0 means that the respective site was not found in both datasets. Red frames indicate the codons selected for toe-printing assay.

**Figure S9.** Toe-printing analysis of BotA2 on mRNAs encoding fMet-Gly-Phe peptide.

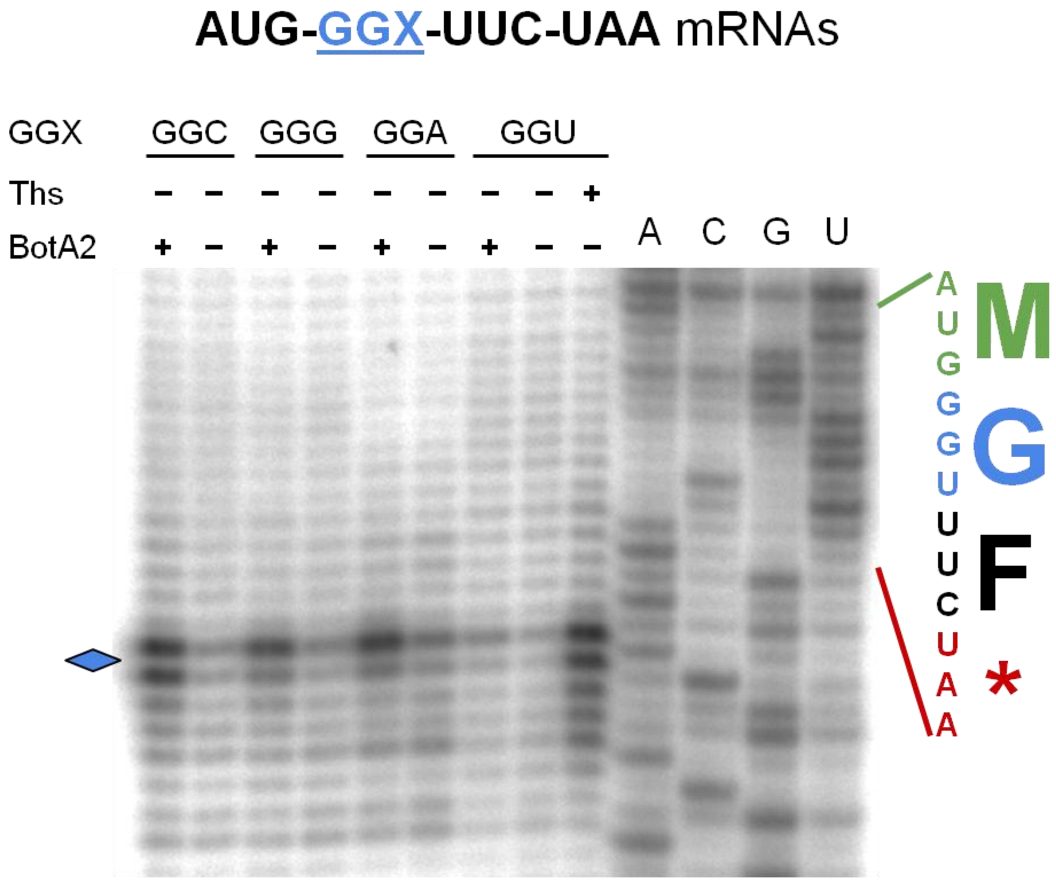

Toe-printing analysis of bottromycin $A_2$ (BotA2) on a set of short mRNA templates encoding fMet-Gly-Phe peptide. All four 3-nt Gly codons were tested to compare the efficiency of BotA2-induced ribosome stalling. Sequences of the corresponding ORF and the encoded amino acids are shown on the right. Asterisk (*) in the translated sequence indicates a stop codon. Rhombus points to the toe-printing bands corresponding to BotA2-induced ribosome stalling. Ths, thiostrepton, was used to indicate the translation start site. All antibiotics were tested at a final concentration of 50 μM.

**Figure S10.** Accumulation of ppGpp in *E. coli* cells upon treatment with BotA2.

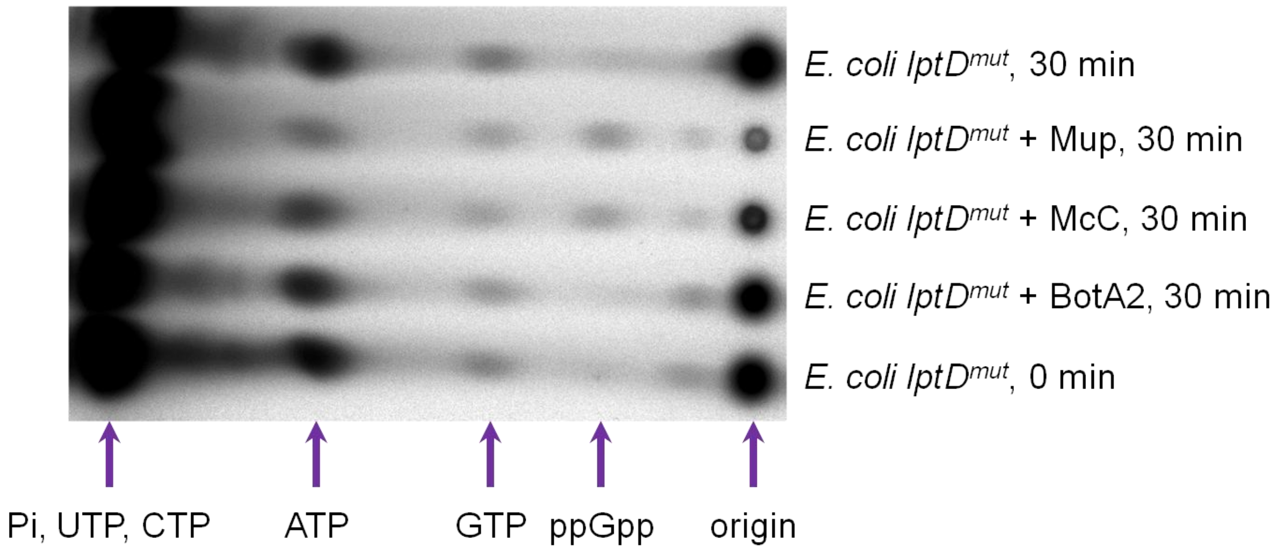

Thin-layer chromatography of [$^{32}$P]-labeled nucleotides isolated from *E. coli lptD$^{mut}$* after 30 min incubation with bottromycin $A_2$ (BotA2), microcin C (McC), or mupirocin (Mup). BotA2 and McC were tested at a final concentration of 20 μM, while Mup was used at a final concentration of 60 μM.

**Figure S11.** Reporter assay for codon misreading by $tRNA^{Glu}$.

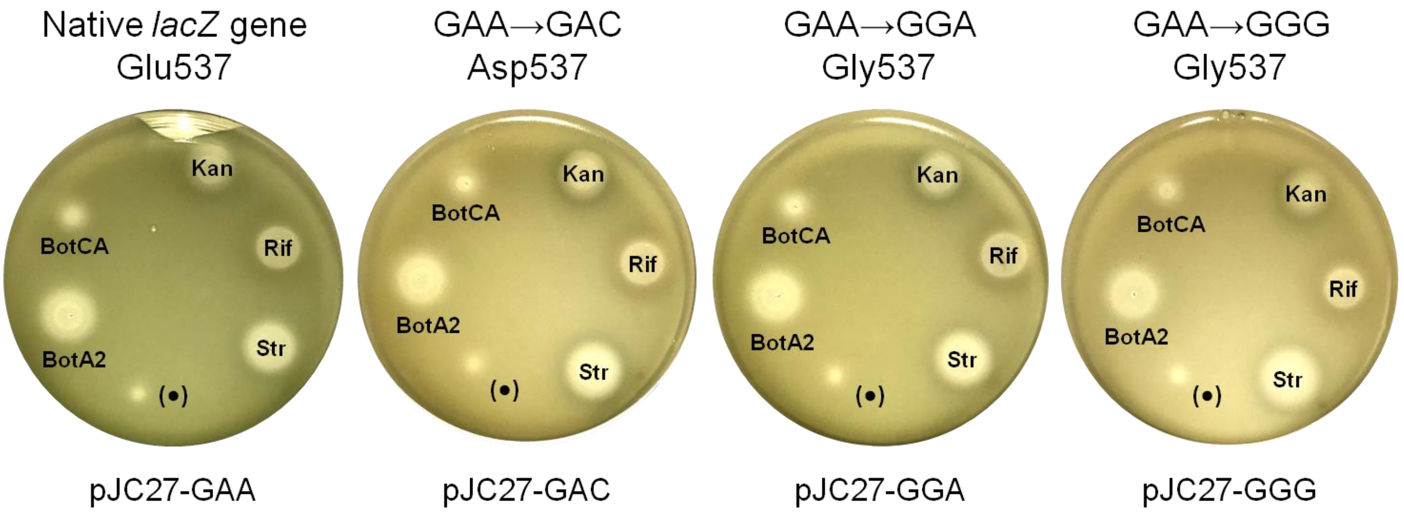

Bottromycin $A_2$ (BotA2) does not cause misreading of the 537$^{th}$ Gly codon of the mutated β-galactosidase gene by Glu-$tRNA^{Glu}$ in *E. coli ΔtolC* cells transformed with reporter systems. The pJC27-GAA construct harbours the native β-galactosidase gene (*lacZ*), in which Glu537 is a catalytic residue essential for the enzymatic degradation of the X-Gal substrate. Constructs pJC27-GAC, pJC27-GGA, and pJC27-GGG harbour mutant variants of the *lacZ* with the 537$^{th}$ GAA codon replaced by GAC, GGA, or GGG, respectively. Miscoding agents, such as kanamycin and streptomycin, reduce the fidelity of translation allowing Glu to be incorporated into the polypeptide sequence at the 537$^{th}$ position of mutated *lacZ* variants. The appearance of blue coloration along the edge of the growth inhibition zone indicates that the antibiotic causes translation errors (mistranslation). The following antibiotics were applied on the surface of an agar plate coated with *E. coli ΔtolC* cells transformed with the indicated pJC27 plasmids: kanamycin (Kan, 5 μg), rifampicin (Rif, 10 μg), streptomycin (Str, 5 μg), bottromycin $A_2$ carboxylic acid (BotCA, 16.2 μg), bottromycin $A_2$ (BotA2, 16.5 μg), and bottromycin $A_2$ (indicated as (•), 0.4 μg). Note that BotCA exhibits smaller growth inhibition zones compared to BotA2, probably due to its impaired penetration into bacterial cells.

**Figure S12.** Codon-anticodon pairing involved in the recognition of Gly codons.

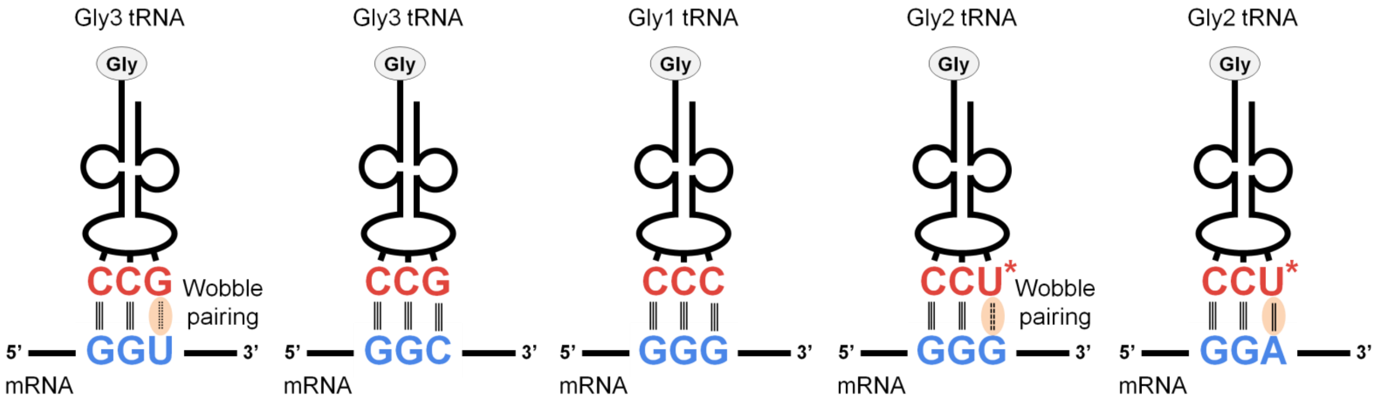

Schematic representation showing the formation of codon-anticodon complexes between Gly codons and cognate Gly-tRNA$^{Gly}$. The canonical Watson–Crick base pairing is shown by vertical lines. Dotted lines represent a wobble base pairing. Dashed lines indicate a wobble base pairing additionally stabilized with a modified uracil (U*). The number of lines reflects the number of hydrogen bonds. Asterisk in the anticodon triplet (U*) indicates a 5-methylaminomethyl-modification of the wobble uridine (mnm$^5$U). Pale orange area highlights weak base pairs at the 3$^{rd}$ poition.

.

**Table S1.** Primers used for the synthesis of DNA templates.

| Primer name | Nucleotide sequence (5’ to 3’) |
|---|---|
| RST1-fwd | AC**TAATACGACTCACTATAGGG**CTTAAGTATAAGGAGGAAAACATATGTATTGGGTAACCTCACGTCAGCCGAATATGCTGAAAATCCATGGCT |
| RST1-rev | GGTTATAATGAATTTTGCTTATTAACGATAGAATTCTATCACTTTTTTTATTATTATTAGGCGCAGTCTTCGAAGCCATGGATTTTCAGC |
| RST3-fwd | AC**TAATACGACTCACTATAGGG**CTTAAGTATAAGGAGGAAAACATATGCATTCTAAATATATATGGGTACTGCGTCAGCCGAATATGAAAGGCTTCG |
| RST3-rev | GGTTATAATGAATTTTGCTTATTAACGATAGAATTCTATCACATTCTTGATTCTTATTAGGCGGTGCAGTCTTCGAAGCCTTTCATATTCG |
| NV1 | GGTTATAATGAATTTTGCTTATTAAC |
| M-AGT-GGC | ATGCA**TAATACGACTCACTATAGGG**CTTAAGTATAAGGAGGAAAACATATGAGTGGCTAAGAGACGGACGAGAGCGGC |
| M-GTC-GGC | ATGCA**TAATACGACTCACTATAGGG**CTTAAGTATAAGGAGGAAAACATATGGTCGGCTAAGAGACGGACGAGAGCGGC |
| M-ATC-GGC | ATGCA**TAATACGACTCACTATAGGG**CTTAAGTATAAGGAGGAAAACATATGATCGGCTAAGAGACGGACGAGAGCGGC |
| M-AAT-GGC | ATGCA**TAATACGACTCACTATAGGG**CTTAAGTATAAGGAGGAAAACATATGAATGGCTAAGAGACGGACGAGAGCGGC |
| M-ACG-GGC | ATGCA**TAATACGACTCACTATAGGG**CTTAAGTATAAGGAGGAAAACATATGACGGGCTAAGAGACGGACGAGAGCGGC |
| M-ATA-GGC | ATGCA**TAATACGACTCACTATAGGG**CTTAAGTATAAGGAGGAAAACATATGATAGGCTAAGAGACGGACGAGAGCGGC |
| M-CTA-GGC | ATGCA**TAATACGACTCACTATAGGG**CTTAAGTATAAGGAGGAAAACATATGCTAGGCTAAGAGACGGACGAGAGCGGC |
| M-CCC-GGC | ATGCA**TAATACGACTCACTATAGGG**CTTAAGTATAAGGAGGAAAACATATGCCCGGCTAAGAGACGGACGAGAGCGGC |
| M-AAA-GGC | ATGCA**TAATACGACTCACTATAGGG**CTTAAGTATAAGGAGGAAAACATATGAAAGGCTAAGAGACGGACGAGAGCGGC |
| M-AGT-GGC-F | ATGCA**TAATACGACTCACTATAGGG**CTTAAGTATAAGGAGGAAAACATATGAGTGGCTTCTAAGAGACGGACGAGAGCGGC |
| M-AGT-GGG-F | ATGCA**TAATACGACTCACTATAGGG**CTTAAGTATAAGGAGGAAAACATATGAGTGGGTTCTAAGAGACGGACGAGAGCGGC |

| | |
|---|---|
| M-AGT-GGA-F | ATGCA**TAATACGACTCACTATAGGG**CTTAAGTATAAGGAGGAAAAC<br>ATATGAGTGGATTCTAAGAGACGGACGAGAGCGGC |
| M-AGT-GGT-F | ATGCA**TAATACGACTCACTATAGGG**CTTAAGTATAAGGAGGAAAAC<br>ATATGAGTGGTTTCTAAGAGACGGACGAGAGCGGC |
| M-GGC-F | ATGCA**TAATACGACTCACTATAGGG**CTTAAGTATAAGGAGGAAAAC<br>ATATGGGCTTCTAAGAGACGGACGAGAGCGGC |
| M-GGG-F | ATGCA**TAATACGACTCACTATAGGG**CTTAAGTATAAGGAGGAAAAC<br>ATATGGGGTTCTAAGAGACGGACGAGAGCGGC |
| M-GGA-F | ATGCA**TAATACGACTCACTATAGGG**CTTAAGTATAAGGAGGAAAAC<br>ATATGGGATTCTAAGAGACGGACGAGAGCGGC |
| M-GGT-F | ATGCA**TAATACGACTCACTATAGGG**CTTAAGTATAAGGAGGAAAAC<br>ATATGGGTTTCTAAGAGACGGACGAGAGCGGC |
| M-AGT | ATGCA**TAATACGACTCACTATAGGG**CTTAAGTATAAGGAGGAAAAC<br>ATATGAGTTAAGAGACGGACGAGAGCGGC |
| M-GGC | ATGCA**TAATACGACTCACTATAGGG**CTTAAGTATAAGGAGGAAAAC<br>ATATGGGCTAAGAGACGGACGAGAGCGGC |
| CER-FAM-R | GCGTTAAGGCTATGTACGGAACAGCTCCTCGCCCTTG |
| NEW_CER_FAM | GCGTTAAGGCTATGTACGCGCCGTCCAGCTCGACCAGG |
| FAMnoFAM | GCGTTAAGGCTATGTAC |
| T7-fwd-1 | ATGCA**TAATACGACTCACTATAGGG** |
| T7-fwd-2 | CGAATT**TAATACGACTCACTATAGG** |
| LP-rev | GCTTGCATGCCTGCAGACGCA |

The sequence of the T7 promoter is marked in bold; the sequence of the NV1 primer used for reverse transcription is underlined; the sequence of the FAMnoFAM primer used for reverse transcription is underlined twice.

**Table S2.** Sequences of DNA templates used in this work

| DNA template | Nucleotide sequence (5’ to 3’) |
|---|---|
| RST1 | AC**TAATACGACTCACTATAGGG**CTTAAGTATAAGGAGGAAAACA<br>TATGTATTGGGTAACCTCACGTCAGCCGAATATGCTGAAAATCCA<br>TGGCTTCGAAGACTGCGCCTAATAATAATAAAAAAAGTGATAGAA<br>TTCTATC**GTTAATAAGCAAAATTCATTATAACC** |
| RST3 | AC**TAATACGACTCACTATAGGG**CTTAAGTATAAGGAGGAAAACA<br>TATGCATTCTAAATATATATGGGTACTGCGTCAGCCGAATATGAA<br>AGGCTTCGAAGACTGCACCGCCTAATAAGAATCAAGAATGTGAT<br>AGAATTCTATC**GTTAATAAGCAAAATTCATTATAACC** |
| M-AGT-GGC<br>(fMet-Ser-Gly) | ATGCA**TAATACGACTCACTATAGGG**CTTAAGTATAAGGAGGAAA<br>ACATATGAGTGGCTAAGAGACGGACGAGAGCGGCCTGGTGAGC<br>AAGGGCGAGGAGCTGTTCC**GTACATAGCCTTAACGC** |
| M-GTC-GGC<br>(fMet-Val-Gly) | ATGCA**TAATACGACTCACTATAGGG**CTTAAGTATAAGGAGGAAA<br>ACATATGGTCGGCTAAGAGACGGACGAGAGCGGCCTGGTGAGC<br>AAGGGCGAGGAGCTGTTCC**GTACATAGCCTTAACGC** |
| M-ATC-GGC<br>(fMet-Ile-Gly) | ATGCA**TAATACGACTCACTATAGGG**CTTAAGTATAAGGAGGAAA<br>ACATATGATCGGCTAAGAGACGGACGAGAGCGGCCTGGTGAGC<br>AAGGGCGAGGAGCTGTTCC**GTACATAGCCTTAACGC** |
| M-AAT-GGC<br>(fMet-Asn-Gly) | ATGCA**TAATACGACTCACTATAGGG**CTTAAGTATAAGGAGGAAA<br>ACATATGAATGGCTAAGAGACGGACGAGAGCGGCCTGGTGAGC<br>AAGGGCGAGGAGCTGTTCC**GTACATAGCCTTAACGC** |
| M-ACG-GGC<br>(fMet-Thr-Gly) | ATGCA**TAATACGACTCACTATAGGG**CTTAAGTATAAGGAGGAAA<br>ACATATGACGGGCTAAGAGACGGACGAGAGCGGCCTGGTGAGC<br>AAGGGCGAGGAGCTGTTCC**GTACATAGCCTTAACGC** |
| M-ATA-GGC<br>(fMet-Ile-Gly) | ATGCA**TAATACGACTCACTATAGGG**CTTAAGTATAAGGAGGAAA<br>ACATATGATAGGCTAAGAGACGGACGAGAGCGGCCTGGTGAGC<br>AAGGGCGAGGAGCTGTTCC**GTACATAGCCTTAACGC** |

| | |
|---|---|
| M-CTA-GGC<br>(fMet-Leu-Gly) | ATGCA**TAATACGACTCACTATAGGG**CTTAAGTATAAGGAGGAAA<br>ACATATGCTAGGCTAAGAGACGGACGAGAGCGGCCTGGTGAGC<br>AAGGGCGAGGAGCTGTTCC**GTACATAGCCTTAACGC** |
| M-CCC-GGC<br>(fMet-Pro-Gly) | ATGCA**TAATACGACTCACTATAGGG**CTTAAGTATAAGGAGGAAA<br>ACATATGCCCGGCTAAGAGACGGACGAGAGCGGCCTGGTGAGC<br>AAGGGCGAGGAGCTGTTCC**GTACATAGCCTTAACGC** |
| M-AAA-GGC<br>(fMet-Lys-Gly) | ATGCA**TAATACGACTCACTATAGGG**CTTAAGTATAAGGAGGAAA<br>ACATATGAAAGGCTAAGAGACGGACGAGAGCGGCCTGGTGAGC<br>AAGGGCGAGGAGCTGTTCC**GTACATAGCCTTAACGC** |
| M-AGT-GGC-F<br>(fMet-Ser-Gly-Phe) | ATGCA**TAATACGACTCACTATAGGG**CTTAAGTATAAGGAGGAAA<br>ACATATGAGTGGCTTCTAAGAGACGGACGAGAGCGGCCTGGTG<br>AGCAAGGGCGAGGAGCTGTTCC**GTACATAGCCTTAACGC** |
| M-AGT-GGG-F<br>(fMet-Ser-Gly-Phe) | ATGCA**TAATACGACTCACTATAGGG**CTTAAGTATAAGGAGGAAA<br>ACATATGAGTGGGTTCTAAGAGACGGACGAGAGCGGCCTGGTG<br>AGCAAGGGCGAGGAGCTGTTCC**GTACATAGCCTTAACGC** |
| M-AGT-GGA-F<br>(fMet-Ser-Gly-Phe) | ATGCA**TAATACGACTCACTATAGGG**CTTAAGTATAAGGAGGAAA<br>ACATATGAGTGGATTCTAAGAGACGGACGAGAGCGGCCTGGTGA<br>GCAAGGGCGAGGAGCTGTTCC**GTACATAGCCTTAACGC** |
| M-AGT-GGT-F<br>(fMet-Ser-Gly-Phe) | ATGCA**TAATACGACTCACTATAGGG**CTTAAGTATAAGGAGGAAA<br>ACATATGAGTGGTTTCTAAGAGACGGACGAGAGCGGCCTGGTGA<br>GCAAGGGCGAGGAGCTGTTCC**GTACATAGCCTTAACGC** |
| M-GGC-F<br>(fMet-Gly-Phe) | ATGCA**TAATACGACTCACTATAGGG**CTTAAGTATAAGGAGGAAA<br>ACATATGGGCTTCTAAGAGACGGACGAGAGCGGCCTGGTGAGC<br>AAGGGCGAGGAGCTGTTCC**GTACATAGCCTTAACGC** |
| M-GGG-F<br>(fMet-Gly-Phe) | ATGCA**TAATACGACTCACTATAGGG**CTTAAGTATAAGGAGGAAA<br>ACATATGGGGTTCTAAGAGACGGACGAGAGCGGCCTGGTGAGC<br>AAGGGCGAGGAGCTGTTCC**GTACATAGCCTTAACGC** |
| M-GGA-F | ATGCA**TAATACGACTCACTATAGGG**CTTAAGTATAAGGAGGAAA |

| | |
|---|---|
| (fMet-Gly-Phe) | ACATATGGGATTCTAAGAGACGGACGAGAGCGGCCTGGTGAGC<br>AAGGGCGAGGAGCTGTTCC**GTACATAGCCTTAACGC** |
| M-GGT-F<br>(fMet-Gly-Phe) | ATGCA**TAATACGACTCACTATAGGG**CTTAAGTATAAGGAGGAAA<br>ACATATGGGTTTCTAAGAGACGGACGAGAGCGGCCTGGTGAGC<br>AAGGGCGAGGAGCTGTTCC**GTACATAGCCTTAACGC** |
| M-AGT<br>(fMet-Ser) | ATGCA**TAATACGACTCACTATAGGG**CTTAAGTATAAGGAGGAAA<br>ACATATGAGTTAAGAGACGGACGAGAGCGGCCTGGTGAGCAAG<br>GGCGAGGAGCTGTTCC**GTACATAGCCTTAACGC** |
| MG<br>(fMet-Gly) | ATGCA**TAATACGACTCACTATAGGG**CTTAAGTATAAGGAGGAAA<br>ACATATGGGCTAAGAGACGGACGAGAGCGGCCTGGTGAGCAAG<br>GGCGAGGAGCTGTTCACCGGGGTGGTGCCCATCCTGGTCGAGC<br>TGGACGGCGC**GTACATAGCCTTAACGC** |
| MF<br>(fMet-Phe-…) | CGAATT**TAATACGACTCACTATAGGG**AATTCAAAAATTTAAAAGT<br>TAACAGGTATACATACTATGTTTACGATTACTACGATCTTCTTCAC<br>TTAATGCGTCTGCAGGCATGCAAGC |
| MV<br>(fMet-Val-…) | CGAATT**TAATACGACTCACTATAGGG**AATTCAAAAATTTAAAAGT<br>TAACAGGTATACATACTATGGTTTTTATTACTACGATCTTCTTCAC<br>TTAATGCGTCTGCAGGCATGCAAGC |

The sequence of the T7 promoter and sequences complementary either to the NV1 primer or to the FAMnoFAM primer are marked in bold. Coding sequences are underlined.

**Table S3.** Parameters of the toe-seq datasets

| | |
|---|---|
| Raw datasets, **before** normalization to controls | |
| Total number of unique mRNAs in all 4 datasets | 57,972 |
| Total number of mRNAs in the BotA2-1 dataset (BotA2 replicate 1) | 56,687 |
| Total number of mRNAs in the BotA2-2 dataset (BotA2 replicate 2) | 56,431 |
| Total number of mRNAs in the Control-1 dataset (Control replicate 1) | 55,835 |
| Total number of mRNAs in the Control-2 dataset (Control replicate 2) | 56,089 |
| Total number of mapped reads in the BotA2-1 dataset (BotA2 replicate 1) | 9,903,509 |
| Total number of mapped reads in the BotA2-2 dataset (BotA2 replicate 2) | 9,360,524 |
| Total number of mapped reads in the Control-1 dataset (Control replicate 1) | 6,148,055 |
| Total number of mapped reads in the Control-2 dataset (Control replicate 2) | 5,923,640 |
| Datasets **after** normalization to controls, **before** filtering the data | |
| Total number of unique mRNAs in 2 datasets (BotA2 replicates 1 and 2) | 56,243 |
| Total number of mRNAs in the BotA2-1 dataset (BotA2 replicate 1) | 55,292 |
| Total number of mRNAs in the BotA2-2 dataset (BotA2 replicate 2) | 55,387 |
| Number of shared mRNAs between two datasets (BotA2 replicates 1 and 2) | 54,436 |
| Number of mRNAs with coincident stalling sites in two datasets (BotA2 replicates 1 and 2) | 13,290 |
| Datasets **after** normalization to controls and **after** filtering the data | |
| Total number of unique mRNAs in 2 datasets (BotA2 replicates 1 and 2) | 24,156 |

| | |
|---|---|
| Total number of mRNAs in the BotA2-1 dataset (BotA2 replicate 1) | 15,995 |
| Total number of mRNAs in the BotA2-2 dataset (BotA2 replicate 2) | 15,694 |
| Number of shared mRNAs between two datasets (BotA2 replicates 1 and 2) | 7,533 |
| Number of mRNAs with coincident stalling sites between two datasets (BotA2 replicates 1 and 2) | 2,854 |

**Table S4.** Antibacterial activity of BotA2 against chloramphenicol-resistant bacterial strains

| | Compound, minimum inhibitory concentration (MIC), µM | | |
|---|---|---|---|
| Strain | BotA2 | BotCA | Chl |
| *E. coli lptD*$^{mut}$ [a] | 6.25 | 25 | 200 |
| *B. subtilis* 168 [b] | 0.78-1.56 | nt | 12.5 |
| *B. subtilis* 168 pHT01-cat [c] | 0.78-1.56 | nt | 200 |
| *B. subtilis* 168 pHT01-cfr [d] | 1.56-3.13 | nt | 100 |

[a] *E. coli lptD*$^{mut}$: chloramphenicol-resistant strain due to the constitutive expression of the *cat* gene, which encodes a chloramphenicol acetyltransferase, CAT.

[b] *B. subtilis* 168: chloramphenicol-susceptible strain.

[c] *B. subtilis* 168 pHT01-cat: chloramphenicol-resistant strain due to the constitutive expression of the *cat* gene, which encodes a chloramphenicol acetyltransferase, CAT.

[d] *B. subtilis* 168 pHT01-cfr: chloramphenicol-resistant strain due to the constitutive expression of the *cfr* gene, which encodes a methyltransferase that modifies A2503 residue of the 23S rRNA.

At least two replicates of MIC measurements were performed.

nt, not tested.

**Table S5.** Characteristics of Gly codons and tRNAs$^{Gly}$ in *Escherichia coli* cells.

| Characteristics of Gly codons in *E. coli* cells | | | | | | |
|---|---|---|---|---|---|---|
| Codon | tRNA$^{Gly}$ | Codon usage in genome, % [2] | Codon usage in transcriptome, % [3] | RSCU, [4] | CEI, [5] | CP, [5] |
| GGU | Gly3 | 2.37 | 3.62 | 2.28 | 14.96 | 698 ± 65 |
| GGC | Gly3 | 2.06 | 2.97 | 1.65 | 4.35 | 514 ± 50 |
| GGG | Gly1<br>Gly2 | 1.23 | 0.92 | 0.04 | -13.39 | 226 ± 19 |
| GGA | Gly2 | 1.36 | 0.63 | 0.02 | -15.15 | 229 ± 19 |

| Characteristics of tRNA$^{Gly}$ in *E. coli* cells | | | | | |
|---|---|---|---|---|---|
| tRNA$^{Gly}$ | Anticodon | Anticodon modifications [6] | Number of tRNA genes [7] | Intracellular concentration of tRNA, μM [3] | Intracellular concentration of free ternary complexes, μM [3] |
| Gly1 | CCC | – | 1 | 4.43 | 3.44-3.51 |
| Gly2 | UCC | mnm$^5$U | 1 | 6.65 | 4.51-4.70 |
| Gly3 | GCC | – | 4 | 24.96 | 14.33-15.38 |

RSCU, relative synonymous codon usage. RSCU values were calculated for highly expressed genes to represent the codon frequency normalized to the expected frequency under the assumption of equal usage of the synonymous codons for Gly amino acid [4].

CEI, codon expression index. CEI shows the level of statistical significance of the correlation between the frequency of a certain codon occurrence in *E. coli* genes and the expression level of the corresponding protein [5]. In other words, CEI reflects if a certain codon has a tendency to be overrepresented in highly expressed genes.

CP, codon productivity [5]. CP shows the average amount of amino acids used by *E. coli* cells for protein production based on a certain codon. CP depends on a codon identity and corresponding tRNA frequencies.

mnm$^5$U, 5-methylaminomethyluridine.
